# A biased allosteric modulator functions as a molecular glue to induce β_2_AR dimerization

**DOI:** 10.1101/2025.10.21.683802

**Authors:** Jiemin Shen, Teja Nikhil Peddada, Konstantin E. Komolov, Francesco De Pascali, Alexander M. Garces, Haoqing Wang, Michael T. Lerch, Jeffrey L. Benovic, Jun Xu, Brian K. Kobilka

## Abstract

Family A G-protein coupled receptors (GPCRs) are typically described as monomers, yet growing evidence suggests they can form dimers with distinct signaling properties^1–3^. The mechanisms and therapeutic potential of such dimerization, however, remain poorly understood. Here, we show that AP-7-168, an optimized derivative of a β-arrestin-biased negative allosteric modulator of the β_2_-adrenergic receptor (β_2_AR) that sustains bronchorelaxation in cell and tissue models^4^, functions as a molecular glue to promote β_2_AR homodimerization. Cryo-EM structures reveal a unique binding mode in which two AP-7-168 molecules pack within a pocket formed by transmembrane helices 3, 4, and 5 of two protomers, stabilizing a dimeric conformation that selectively prevents β-arrestin coupling. In cells, AP-7-168 robustly induces β_2_AR dimerization and drives enlarged nanocluster formation. Combined with extensive functional studies, our findings unveil a novel allosteric mechanism by which a small molecule biases β_2_AR signaling through dimerization, highlighting ligand-induced dimerization as a strategy for GPCR modulation.

## Introduction

G-protein coupled receptors (GPCRs) represent the largest superfamily of transmembrane (TM) proteins, exhibiting complex signaling behaviors. Depending on the ligand, GPCRs can interact with G-proteins or bind to effector proteins such as arrestins and kinases, mediating G-protein-independent pathways and giving rise to diverse pharmacological and physiological responses^5,6^. Over 800 human GPCRs have been identified and systematically classified into five families based on their sequence features and structural characteristics^7^: i) family A Rhodopsin-like, ii) family B1 Secretin-like, iii) family B2 Adhesion-like, iv) family C Glutamate-like, and v) family F Frizzled-like receptors. Although family C receptors are known to be obligate dimers^8^, mediated primarily by their large Venus flytrap extracellular domains (ECDs), dimerization of other families is not well described.

Over the past few decades, many studies suggest that family A receptors also possess the ability to form dimers^9–11^. Recent cryo-EM studies on rhodopsin^1^, apelin receptor (APJR)^2^, and GPR3^3^ provide direct evidence for the dimerization of these family A GPCRs. However, unlike family C receptors, family A receptors do not have a significant ECD and therefore rely on interactions between their TM domains to undergo dimerization^12^. Despite the various studies on family A dimers, their physiological relevance is still debated. The development of ligands that stabilize family A receptor dimers remains elusive^13,14^, and it is difficult to reliably reproduce these dimers and to comprehensively study their functional, biochemical, and biophysical properties.

We previously identified a class of negative allosteric modulators (NAMs) selective for the β_2_-adrenergic receptor (β_2_AR), a family A GPCR, that exhibited biased inhibition of β-arrestin recruitment with minimal impact on stimulatory G protein (Gs)-mediated cAMP production in cell-based assays^4^. Through comprehensive structure–activity relationship (SAR) studies, we developed AP-7-168 (hereafter referred to as AP), a compound that has high potency and biased effects. Activation of β_2_AR upon coupling with Gs triggers the downstream cAMP signaling pathway, leading to many physiological effects including the relaxation of airway smooth muscle, which is the core mechanism underlying the use of β_2_AR agonists in treating airway diseases such as asthma and chronic obstructive pulmonary disease. However, upon prolonged agonist stimulation, β_2_AR undergoes desensitization via arrestin-mediated inhibition of Gs coupling and receptor endocytosis, resulting in a diminished bronchodilatory effect. Our *ex vivo* tissue assays demonstrated that AP effectively maintained agonist-induced β_2_AR activation and the subsequent bronchodilation while significantly delaying receptor desensitization^4^, highlighting its potential clinical value. Despite these promising findings, the molecular mechanism by which AP modulates β_2_AR function remains unclear, posing a significant obstacle to the further development and optimization of AP as a therapeutic agent.

In this study, using cryogenic electron microscopy (cryo-EM) combined with various biochemical and biophysical approaches, we demonstrate that AP functions as a molecular glue by stabilizing a homodimer complex of β_2_AR, which in turn alters the downstream signaling behaviors. These findings reveal a novel allosteric modulation mechanism in β_2_AR that may have broad implications in drug development for family A GPCRs.

## Results

### Structure determination of AP-bound β_2_AR

As depicted in Figure 1A, AP and its parent compound difluorophenyl quinazoline (DFPQ)^4^ have the pharmacophore characterized by a quinazoline ring with 2- and 4-amino substitutions. AP features a 3,4-difluorophenyl group at the 2-amino position, a cyclohexane substituent at the 4-amino position, and a bromine atom at the 6-position of the quinazoline ring. AP exhibited a remarkable biased NAM effect in β_2_AR-expressing cells, being approximately 1,000-fold more effective at inhibiting β-arrestin2 recruitment versus cyclic adenosine monophosphate (cAMP) production when stimulated with the β-agonist isoproterenol (Iso) (Figure 1B). To elucidate the molecular mechanism underlying this biased inhibition, we sought to determine the structure of β_2_AR bound to AP.

**Figure 1.**
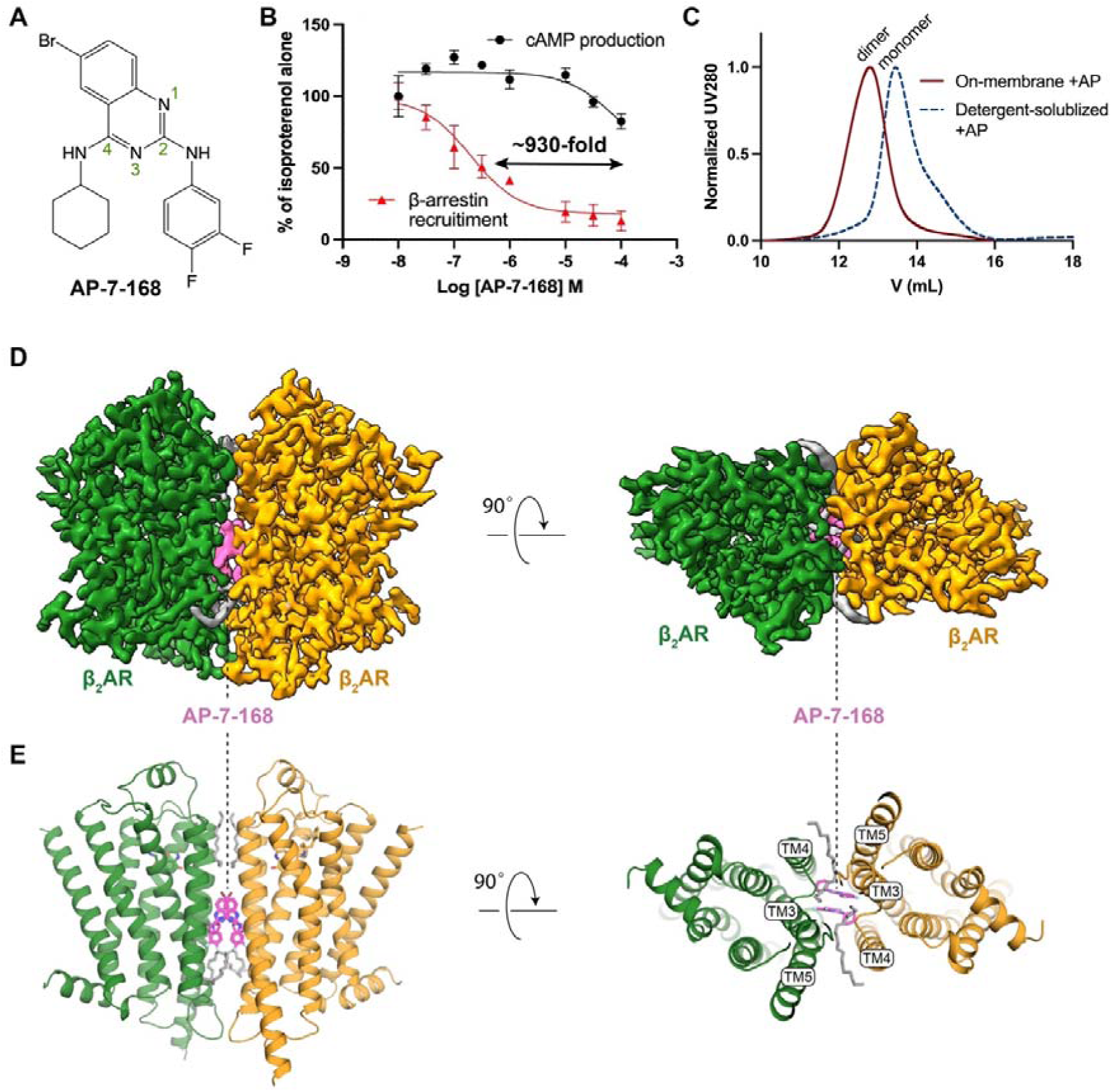
Cryo-EM structures reveal that AP, a β-arrestin–biased NAM, mediates β_2_AR dimerization. (**A**) Chemical structure of AP-7-168 (AP). Atom numbering on the quinazoline ring is indicated. (**B**) Dose-dependent effects of AP on β-arrestin recruitment (red) and cAMP production (black) in HEK cells stimulated with 1 µM isoproterenol (Iso). AP exhibits ∼930-fold lower IC_50_ for β-arrestin recruitment. Data are normalized to Iso-alone controls and presented as mean values with error bars representing the standard error of the mean (SEM), n = 3. (**C**) Size-exclusion chromatography (SEC) profiles of β_2_AR treated with AP on the cell membrane (red) versus detergent-solubilized β_2_AR with subsequent AP addition (black). (**D**) Cryo-EM map of AP-bound β_2_AR dimer reconstituted in lipid nanodiscs. Densities for two protomers are colored in green and orange, densities for AP are colored in magenta and densities for the lipids are colored in gray. (**E**) Overall structure of the AP-bound β_2_AR dimer.

We initially prepared the cryo-EM sample using detergent-solubilized and size-exclusion chromatography (SEC)-purified β_2_AR incubated with AP in the presence of carazolol, a β-antagonist. Given the small size (< 50 kDa) of a monomeric receptor, we included nanobody 60 (Nb60), which is specific to an inactive conformation of β_2_AR^15^, for steering particle alignment in cryo-EM data processing. Unexpectedly, the extracted particles from the dataset revealed two predominant 2D class averages, with approximately 70% of monomers and 30% of dimers (Figure S1A). Subsequent 3D classification and reconstruction showed the dimer consists of two parallel protomers with a symmetric TM3, TM4, and TM5 interface (Figure S1). Owing to the larger size and the C2 symmetry of the dimer, we obtained a cryo-EM map at a resolution of 2.9 Å (Figure S1A). The map revealed two distinct AP densities located at the dimer interface near the center of the membrane (Figure 1D and S1A). This finding suggests that AP may promote β_2_AR dimerization. We also managed to reconstruct a 4.3 Å resolution map for monomeric β_2_AR bound to Nb60 in the same dataset (Figure S1A). Despite the modest resolution, a distinct non-protein density, comparably strong to that of TM helices, was observed near the AP binding site identified in the dimer structure (Figure S2A and S2B). This indicates that AP can bind to monomeric β_2_AR at a similar position as in the dimer. Surface plasmon resonance (SPR) measurements further confirmed the direct binding of AP to the monomeric receptor with a *K*_D_ of approximately 30 μM (Figure

S2C).

Encouraged by the initial dimeric structure in detergent, we sought to produce a homogeneous dimer sample and solve its structure in a native-like environment. Although incubation of AP with purified monomeric β_2_AR in detergent did not result in efficient dimerization as assessed by SEC (Figure 1C), we achieved almost complete dimerization when β_2_AR was pretreated with AP in a membrane environment before detergent extraction (Figure 1C and Methods). This highlights the critical role of a continuous fluid membrane in facilitating TM contacts mediated by AP and maintained in detergent after solubilization. We reconstituted the dimer into lipid nanodiscs and solved its structure in complex with an agonist, BI-167107 (BI), without nanobodies (Figure S1B). Cryo-EM analysis confirmed the presence of predominantly dimeric species, with no monomer detected in 2D classification (Figure S1B). The final cryo-EM map achieved a resolution of 2.5 Å, with a local resolution of ∼2.2 Å near the AP binding site (Figure 1D, 1E, and S1B).

### Structural basis of AP binding

The high-resolution map enabled precise modeling of two AP molecules in the dimer, revealing a novel ligand binding mode (Figure 1E and S3). The two AP molecules are packed against each other through π–π and van der Waals interactions and fit in an allosteric pocket formed by TM3, TM4, and TM5 of two β_2_AR protomers (Figure 1E and 2A–2C). The two AP molecules also form extensive hydrophobic and van der Waals interactions with a series of non-polar residues in TM3 (C125^3.44^, V126^3.45^, V129^3.48^), TM4 (M156^4.48^, M157^4.49^), and TM5 (V206^5.45^, V210^5.49^, I214^5.53^) (Figure 2A–2C). In addition, the 2-amino group in AP forms hydrogen bond interactions with E122^3.41^ in TM3 (Figure 2A). These specific interactions in our structural model rationalize previously reported SAR and mutagenesis data^4^. For example, removing the bromine atom reduced AP’s activity, likely because of the loss of interactions with V206^5.45^ (Figure 2A). The difluorophenyl substitution forms optimal contacts with V126^3.45^ and V129^3.48^ (Figure 2C), whereas larger halogen atoms reduce ligand potency^4^. Similarly, V129^3.48^L or swapping TM3 with β_1_-adrenergic receptor (β_1_AR), where isoleucine replaces the valine residue, introduces steric clashes with the difluorophenyl group, therefore significantly reducing the potency of AP. Moreover, the E122^3.41^W mutation also leads to almost complete loss of response to AP^4^.

**Figure 2.**
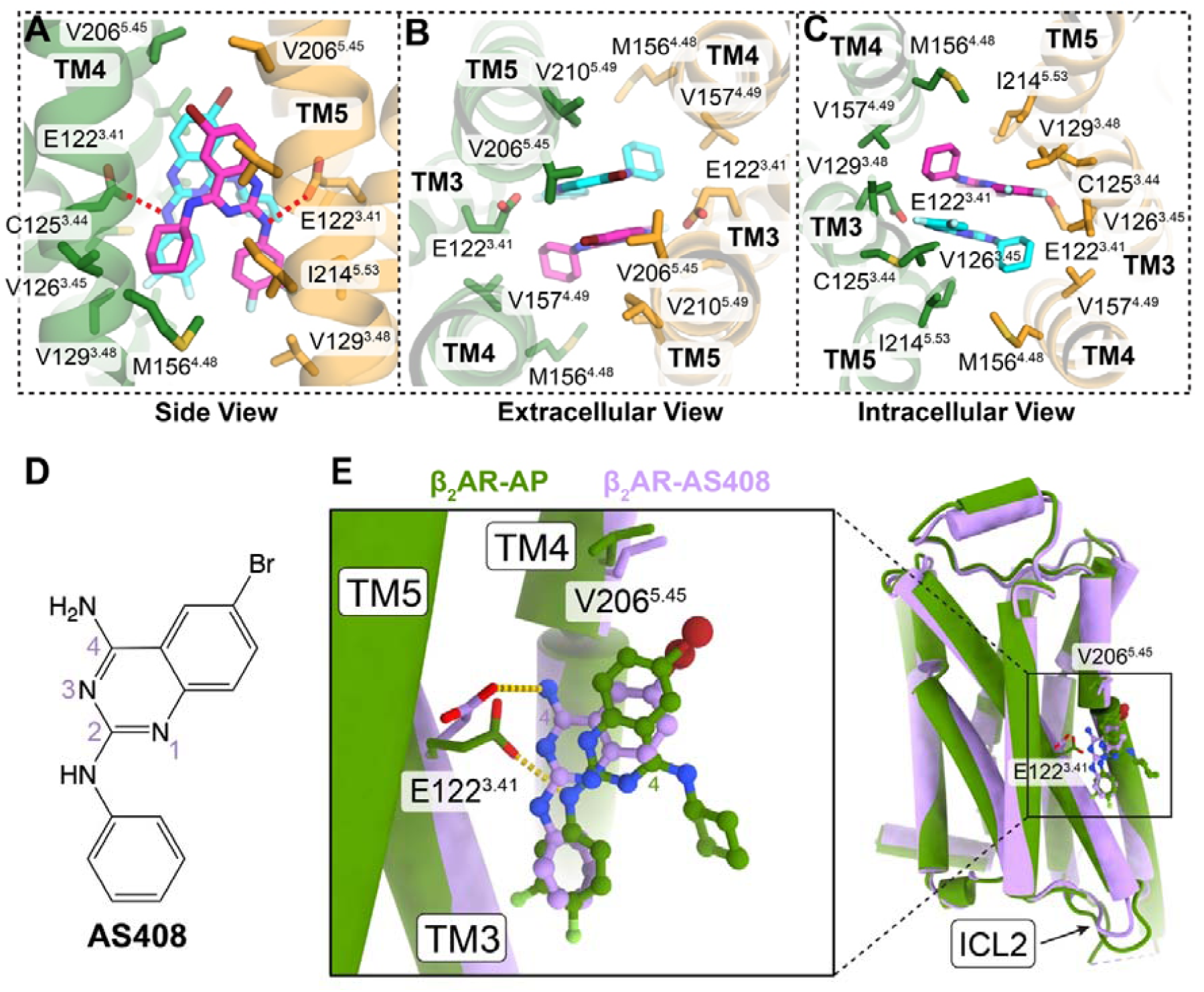
Structural analysis of AP binding site. (**A–C**) Enlarged AP binding pocket from side view (**A**), extracellular view (**B**) and intracellular view (**C**). The two AP molecules are shown in magenta and cyan respectively. Residues within 4 Å of AP were shown in sticks. Red dashed lines represent hydrogen bond interactions. (**D**) Chemical structure of the unbiased NAM, AS408. (**E**) Structural comparison of one protomer from the AP-bound β_2_AR dimer (green) with monomeric β_2_AR bound to AS408 (purple, PDB ID: 6OBA). Enlarged view of the ligand binding pocket is shown on the left. C4 atoms of the quinazoline ring are labeled to highlight the 180° flip between AP and AS408.

Interestingly, the binding site of AP on β_2_AR closely overlaps with a previously identified binding site for an unbiased NAM, AS408^16^ (Figure 2D and 2E). Structural alignment of one of the β_2_AR protomers in the AP-bound dimer with the β_2_AR-AS408 complex (PDB ID: 6OBA) reveals a highly similar overall conformation, with an overall Cα root mean square deviation (RMSD) of only 0.53 Å (Figure 2E). Both AP and AS408 possess a bromide substitution at the C6 position of the quinazoline ring (Figure 2D and 2E), and this bromide is positioned in close proximity to V206^5.46^ in both structures (Figure 2E). However, despite the similarities in the quinazoline ring and the bromide position, there are notable differences in their binding orientations.

The quinazoline rings of AP and AS408 are nearly parallel, but they are flipped 180 degrees relative to each other (Figure 2E). This difference in orientation is likely due to the absence of a bulky substitution at the 4-amino position in AS408, in contrast to AP. Consequently, in the AS408-bound structure, the 4-amino group of AS408 forms a hydrogen bond with E122^3.41^, while in the AP-bound structure, it is the 2-amino group of AP that interacts with E122^3.41^ (Figure 2E).

E122^3.41^ has been proposed to be a critical conformational hub in β_2_AR^16^. It serves as a key link that connects the extracellular orthosteric ligand-binding pocket to the intracellular transducer-coupling interface. The shift in hydrogen bonding partners, from the 4-amino group of AS408 to the 2-amino group of AP, stabilizes E122^3.41^ in a distinct conformation (Figure 2E). This unique conformation of E122^3.41^ may have significant implications for modulating the interactions of β_2_AR with different intracellular transducers, such as Gs and β-arrestins. Moreover, the cyclohexane group in AP, which is not present in AS408, plays a crucial role in ligand packing and hydrophobic contacts with the adjacent β_2_AR protomer. These differences may explain why AS408 lacks the ability to promote receptor dimerization and the biased NAM effect of AP.

### AP-mediated **β**_2_AR dimer interface

The AP-mediated β_2_AR dimer interface is primarily built by TM3, TM4, and TM5, along with the loop regions ECL2 and ICL2 (Figure 3A). In addition to the AP-mediated interactions around the central binding pocket (Figure 2A–2C), there are extensive intermolecular interactions between the two β_2_AR protomers from both extracellular and intracellular sides. As shown in Figure 3, in the extracellular side, L167^4.59^ and M171^4.63^ in TM4 and W173 in ECL2 of one protomer form extensive hydrophobic and van der Waals interactions with Q197^5.36^, A198^5.37^, I201^5.40^, and I205^5.44^ in TM5 of the other protomer. On the intracellular side, K149^4.41^ in TM4 forms a hydrogen-bonding interaction with the conserved Y132^3.51^ (of the DRY motif) in TM3 of the adjacent protomer. In addition, F133^3.52^ in TM3 and F139^34.51^, K140^34.51^, L144^34.56^, and L145^34.57^ in the ICL2 loop create an extensive hydrophobic network between the two protomers. Moreover, we observed clear lipid-like densities, running almost parallel to the membrane plane, bridging hydrophobic contacts between the two protomers (Figure 3), which further stabilize the dimer conformation of β_2_AR.

**Figure 3.**
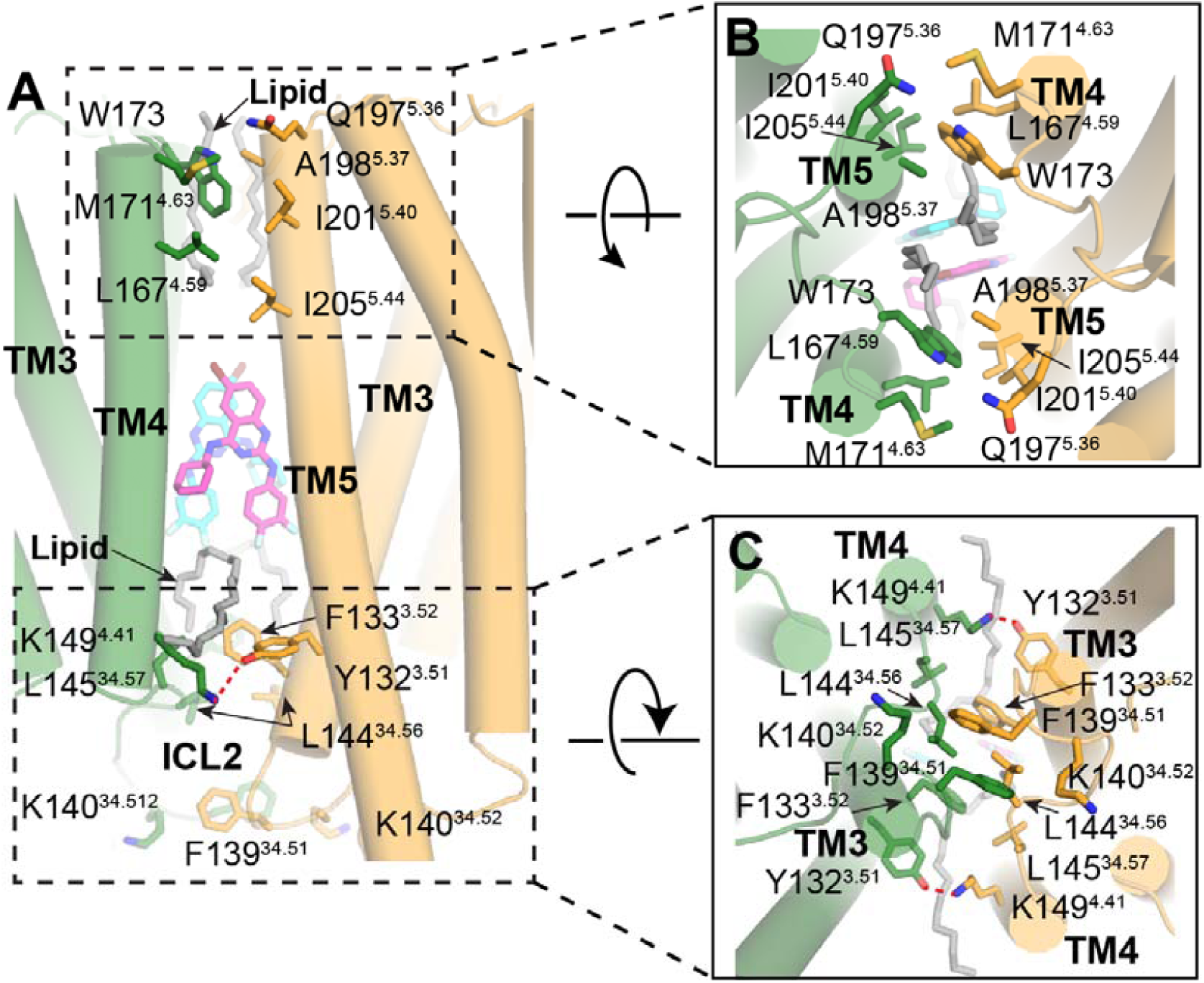
AP-mediated β_2_AR dimer interface. **(A-C)** Detailed interactions between two protomers in the dimer interface from different views: (**A**) side view, (**B**) extracellular view and (**C**) intracellular view. Residues that are within 4 Å between the two protomers are shown as sticks. Red dashed lines represent hydrogen bond interactions. The lipid molecules in the dimer interface are shown in gray sticks.

The β_2_AR dimer interface captured in our cryo-EM structures differs from the potential interface observed in the lipid cubic phase (LCP) crystal structure of β_2_AR^17^ (Figure S4A). The AP-stabilized β_2_AR dimer interface also contrasts with those in the APJR dimers^2,12,18,19^ (Figure S4B), rhodopsin dimer^1^ (Figure S4C), and GPR3 dimer^3,20,21^ (Figure S4D), while it is similar to a potential β_1_AR dimer interface in the LCP crystal structure^22^ (Figure S4E). While previous biophysical studies have shown that the β_2_AR could form dimers and higher-order oligomers by itself^10,23^, it is possible that the new interface in our structures represents a native yet transient one for β_2_AR.

### Validation of AP-mediated **β**_2_AR dimerization in liposomes and in cells

The above structures of AP-bound β_2_AR dimer were resolved in detergent micelles or nanodiscs. To validate whether the AP-mediated β_2_AR dimer adopts the same structural arrangement in a continuous membrane, we applied single-molecule fluorescence resonance energy transfer (smFRET) microscopy to probe its conformation in liposomes. As shown in Figure 4A, the TM helices in the dimer structure resolved by cryo-EM form a parallelogram-like arrangement when viewed perpendicular to the membrane plane, with two diagonals of different lengths. Fluorescent donor and acceptor dyes labeled on residues near different vertices should yield distinct inter-dye distances and corresponding FRET values. To label each protomer in a site-specific manner, single-cysteine mutations were introduced into a minimal cysteine construct of β_2_AR^24^ and “clicked” with maleimide-conjugated fluorescent dyes. Stochastic labeling is expected to result in half of the doubly labeled dimers having one donor and one acceptor dye. The labeled proteins were surface-immobilized and imaged using objective-based total internal reflection fluorescence (TIRF) microscopy. TM5-labeled samples exhibited a homogeneous Gaussian distribution of FRET values centered at ∼0.9, while helix 8 (H8)-labeled samples showed a distribution centered at ∼0.5 (Figure 4A), with the higher FRET consistent with a shorter distance between the fluorophore pairs. These results are consistent with the cryo-EM structure and indicate the conformational stability of the dimer. To assess long-term stability, we measured ensemble FRET changes using dye-labeled samples in detergent (Figure S5A). When dimers were diluted into AP-free buffer, the relative FRET values remained nearly unchanged over 24 hours (Figure S5B and S5C). Moreover, incubation with Gs or activated β-arrestin could not dissociate the dimer (Figure S5D).

**Figure 4.**
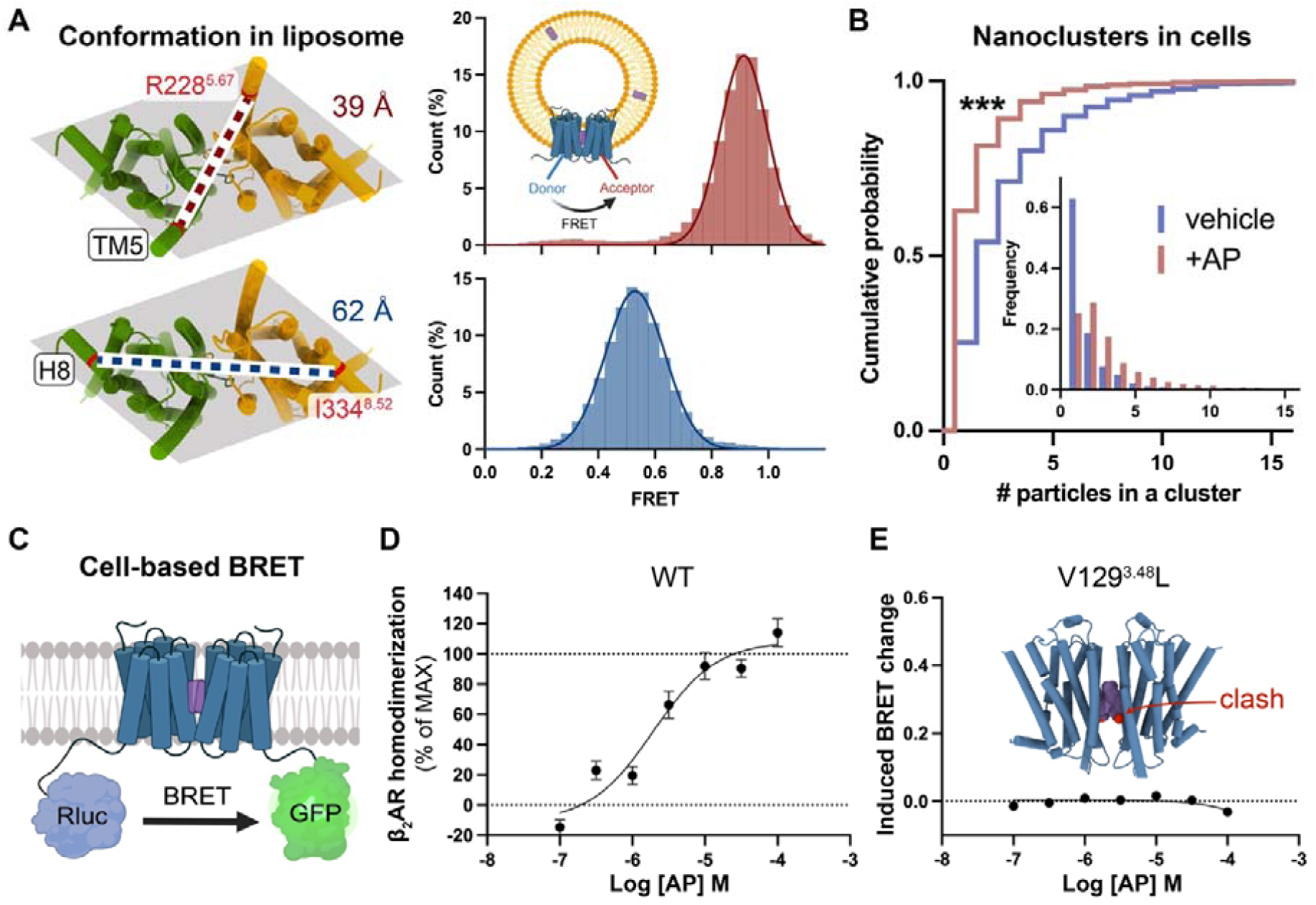
AP promotes β_2_AR dimerization in continuous membranes and in cells. (**A**) smFRET assay showing the conformation of AP-bound β_2_AR dimer in proteoliposomes. Shaded parallelograms indicate the geometry of the dimer viewed from the intracellular side. Dye labeling sites (red) on TM5 (top) and H8 (bottom) and their respective inter-protomer distances (dotted lines) are indicated on the structure, with corresponding FRET population histograms shown on the right. Inset: schematic of β_2_AR dimer in a liposome with inter-protomer FRET; purple rectangles represent AP molecules. (**B**) Cumulative probability plots of β_2_AR cluster size on the cell membrane. Inset: frequency distributions of cluster size. Blue, vehicle; red, treatment with 10 µM AP. ***, p < 0.001. (**C**) Schematic of the cell-based BRET assay for receptor dimerization. (**D**) Dose-response curve of AP-promoted β_2_AR dimerization in HEK cells. (**E**) The V129^3.48^L mutation, which introduces steric clashes with AP, abolishes β_2_AR dimerization.

The cryo-EM structure and biophysical data presented above provide strong evidence that AP promotes β_2_AR dimerization; however, it remains unclear whether this effect occurs in cells. To further address this, we employed two techniques to assess β_2_AR dimerization in cells. First, we analyzed the size of β_2_AR nanoclusters with or without AP treatment. GPCRs are known to form mesoscale nanoclusters in cells, which act as signaling hubs to locally concentrate effectors and enhance signaling efficiency^25,26^. Molecular mechanisms underlying nanoclustering are poorly understood but likely involve weak but specific interactions among receptors (Figure S6A). We hypothesized that AP stabilizes the dimer as a single unit that has increased interaction sites, facilitating prolonged contacts and the assembly of larger receptor clusters (Figure S6A). To visualize nanocluster distribution on cell membranes, we labeled β_2_AR with a gold nanoparticle (AuNP)-conjugated antibody and prepared cryo-EM grids with cytoplasmic content removed via unroofing. As shown in Figure S6B-C, cryo-EM imaging revealed distinct AuNP clusters rather than random scatters. Statistics of AuNPs in each cluster demonstrated that AP significantly increased nanocluster size (Figure 4B) as a result of AP-induced dimerization of β_2_AR. The resulting enlarged nanoclusters may amplify signaling outcomes compared to isolated monomers or dimers combined.

We also performed cell-based Bioluminescence Resonance Energy Transfer (BRET) assays to quantify β_2_AR dimerization by measuring distance-dependent BRET signals between Rluc-tagged and GFP-tagged receptors (Figure 4C). Cells co-expressing differently tagged β_2_AR exhibited AP-dependent dimerization with a pEC_50_ of 5.93 ± 0.18, n = 4 (Figure 4D). Control experiments confirmed the specificity of this effect, showing no fluorescence interference from AP, no dimerization induced by Iso alone, and no expression-level-dependent self-homodimerization (Figure S7A–S7C). To further validate dimerization specificity, we tested a β_2_AR mutant (V129^3.48^L) that disrupts the AP binding pocket due to steric clashes. As expected, this mutation completely abolished AP-mediated dimerization (Figure 4E). Similar effects were observed when swapping TM3 from β_1_AR into β_2_AR (Figure S7D) or assessing AP-mediated β_1_AR dimerization (Figure S7E). These results collectively demonstrate that AP robustly promotes β_2_AR dimerization in native cellular environments.

### Higher-order oligomerization of **β**_2_AR

Apart from the dimer species in the cryo-EM dataset, a subset of 2D classes displayed top views of higher-order oligomers, including tetramers and hexamers (Figure S8A). We reconstructed a low-resolution map of the tetramer, revealing a dimer-of-dimers organization (Figure S8B). Docking of two dimer structures into the map shows symmetric interactions between the dimers, mediated by the intracellular regions of TM5–TM6 and TM1–ICL1 (Figure S8C). Mass photometry of diluted dimer samples in detergent confirmed the presence of a significant fraction of tetramer in solution (Figure S8D–S8F). Consistent with these findings, immunogold imaging of cell membranes showed enlarged nanoclusters upon AP treatment (Figure 4B and S6), supporting the notion that the AP-induced dimer serves as a structural unit for higher-order oligomerization. Together, these results indicate that AP-bound β_2_AR can assemble into higher-order oligomers, providing a structural basis for receptor nanocluster formation in cells.

### The **β**_2_AR dimer functions as a biased signaling species

To understand the structural basis of the biased NAM activity of AP, we first compared our dimer structure with previously resolved monomeric β_2_AR structures. One protomer in the dimer structure aligns well with the previous crystal structure of β_2_AR in an inactive conformation (PDB ID: 2RH1), with an overall Cα RMSD of 0.53 Å (Figure 5A). Notably, the structure of the dimer in nanodiscs, resolved without nanobodies or engineered fusions, exhibits a distinct ICL2 conformation from the crystal structure with an Cα RMSD of 1.5 Å (Figure 5A). F139^34.51^ on ICL2 participates in dimer contact with the same residue on the opposing protomer (Figure 3), resulting in an outward shift away from the central cavity involved in G-protein engagement (Figure 5B). This shift is accompanied by other conformational changes in ICL2 (Figure 5B and S3B): Y141^34.53^ forms hydrogen bond and π–cation interactions with R131^3.50^ in TM3, which normally forms a hydrogen bond with H269^6.31^ in TM6; K263^6.25^ at the intracellular end of TM6 appears to replace K140^34.52^ on ICL2 in neutralizing the helical dipole of TM3 (Figure S3B). These local conformational rearrangements may alter the interplay between β_2_AR and its transducers.

**Figure 5.**
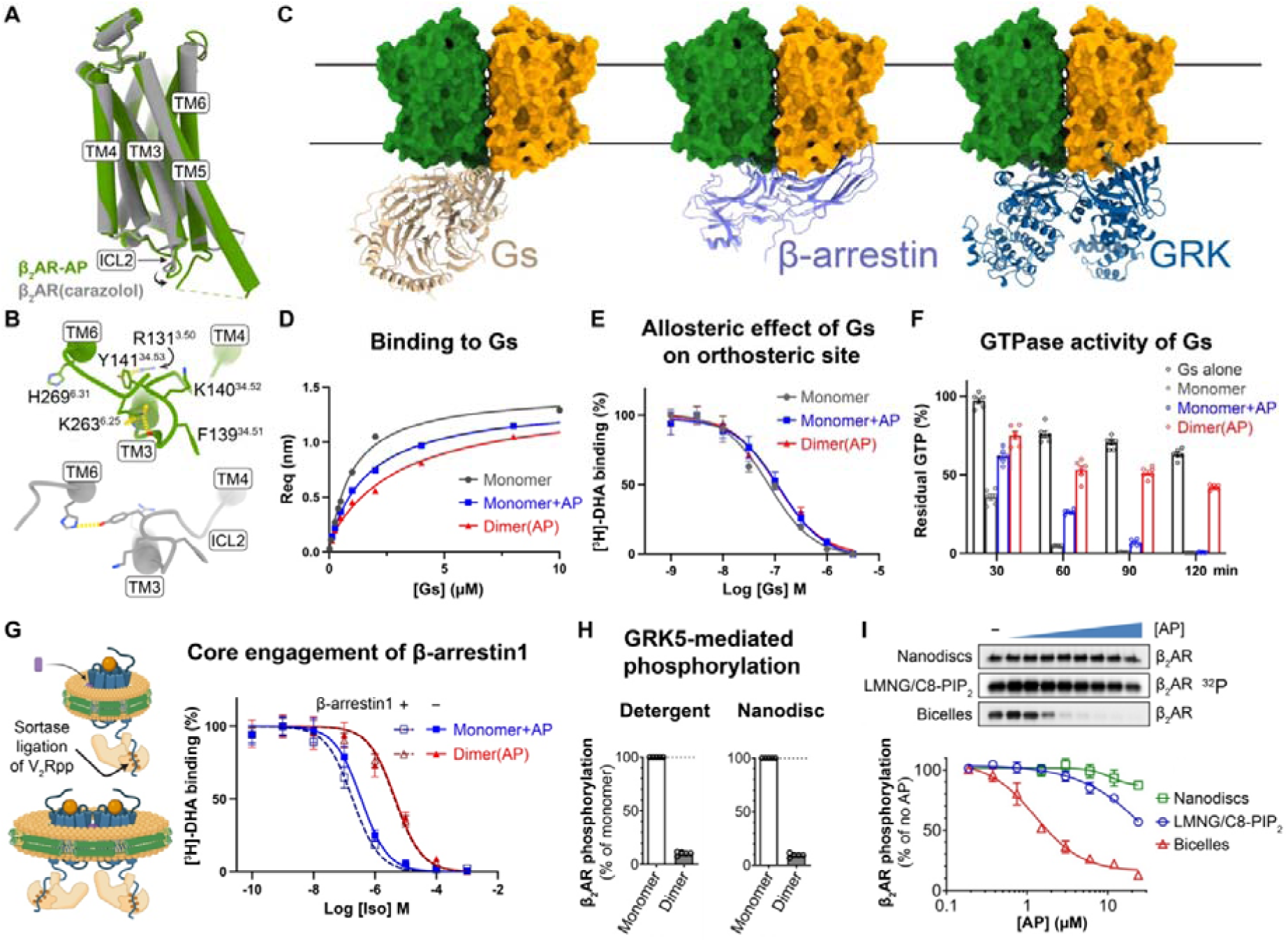
AP-bound β_2_AR dimer is a species with biased activity. (A) Overall structure comparison of the AP-bound β_2_AR dimer (one protomer, green) with the carazolol-bound β_2_AR crystal structure (2RH1, gray) reveals an inactive state conformation. **(B)** Comparison of the ICL2 conformation between the AP-bound β_2_AR (green) and the carazolol-bound inactive β_2_AR (gray). Yellow dashed lines represent hydrogen bond interactions. (**C**) Structural models of β_2_AR dimer (green and orange) aligned with known transducer-bound GPCR structures: Gs (wheat), β-arrestin (lavender), and GRK (dark blue). Models were generated by aligning monomeric effector-bound structures to one protomer of the AP-bound β_2_AR dimer. Original structures: β_2_AR–Gs (PDB: 3SN6), β_1_AR–β-arrestin1 (PDB: 6TKO), NTR1–GRK2 (PDB: 8JPB). (**D**) Steady-state binding curves of GDP-bound Gs to β_2_AR measured by BLI. (**E**) Dose-dependent negative allosteric effect of Gs binding on antagonist binding to the orthosteric site of β_2_AR. Measurements were performed on β_2_AR monomers and dimers reconstituted in lipid nanodiscs. (**F**) GTP turnover assays show the different NAM effects of AP on the GEF activity of monomeric and dimeric β_2_AR toward Gs. Reactions were quenched at indicated time points and residual amounts of GTP were quantified. (**G**) Schematic of monomeric or dimeric β_2_AR–V_2_Rpp in nanodiscs with AP, showing β-arrestin tail engagement but no core interactions. Right: Competition binding of [³H]-DHA with agonist Iso in monomer (blue) or dimer (red) in the presence (dash line) or absence (solid line) of β-arrestin1. 10 μM AP was included. (**H**) Radiometric kinase assay showing phosphorylation of β_2_AR monomers and dimers solubilized in LMNG detergent or reconstituted into nanodiscs. (**I**) Dose-dependent effect of AP on β_2_AR phosphorylation. β_2_AR monomers were solubilized in MNG detergent or reconstituted into bicelles or nanodiscs and treated with AP for 30 min before the reaction. Top: ^32^P autoradiography of β_2_AR. Data are shown as mean with SEM; n = 3 in (**E**), n = 6 in (**F**), n = 3 in (**G**), and n = 5 in (**H**) and (**I**).

Structural analysis of β_2_AR bound to its transducers provided additional insights into how the dimer achieves biased inhibition. Alignment of β_2_AR–Gs^27^, β_1_AR–β-arrestin1^28^, and neurotensin receptor type 1 (NTSR1)–G protein-coupled receptor kinase 2 (GRK2)^29^ structures with one protomer in the dimer (Figure 5C) revealed significant steric clashes between β-arrestin1 or GRK2 and the intracellular side of the other protomer, preventing their engagement with the receptor. In contrast, docked Gs appeared to be accommodated. However, dimerized β_2_AR adopts an inactive-like conformation. Structural comparison with the Gs-bound receptor (Figure S9A) revealed that local conformational changes breaking dimer symmetry are needed to allow for the loop-to-helix transition in ICL2 (Figure S9B and S9C), necessary for engagement with Gs in the canonical state, while the outward movements of TM5 and TM6 are not restricted by the dimer contacts. Together, these data suggest that dimerization of the β_2_AR contributes to the biased NAM effect of AP.

To further confirm the functional consequences of the AP-induced dimerization, we conducted a series of biochemical assays to investigate the *in vitro* activities of the β_2_AR monomer and dimer in isolation. Unlike cell-based assays, where β_2_AR may exist as a mixture of monomers and dimers after AP treatment, the homogeneous and stable dimer sample that we obtained from purification enabled us to characterize the functions of dimer and monomer individually. First, we demonstrated that the dimer could form a stable complex with nucleotide-free Gs. This complex eluted as a single peak in SEC (Figure S10A), and the peak fractions contained the bands for Gs heterotrimer on SDS-PAGE (Figure S10B). Estimation of the relative ratios of β_2_AR to Gα on gels indicated a 2:1 stoichiometry in the dimer– Gs complex, consistent with a docking model in which two Gs molecules cannot simultaneously engage the dimer (Figure 5C). Preliminary cryo-EM analysis of the dimer–Gs complex shows 2D class averages with one Gs density associated with an oval-shaped micelle housing one dimer (Figure S10C).

We then used bio-layer interferometry (BLI) to measure the binding affinities and kinetics of guanosine diphosphate (GDP)-bound Gs to monomeric or dimeric β_2_AR, with and without AP (Figure 5D and S10D–S10F). All samples exhibited μM-level affinity for Gs(GDP) (Table S2). Despite adopting an inactive conformation in the structural models, the β_2_AR dimer retains the ability to engage Gs, suggesting that the dimeric β_2_AR can adopt a Gs-favorable conformation. The dimer showed a modest (∼3-fold) decrease in affinity for Gs (dissociation constant, *K*_D_ = 3.11 μM) compared to the monomer (*K*_D_ = 0.96 μM). AP at 10 μM caused a slight reduction in the monomer–Gs binding affinity (*K*_D_ = 1.7 μM). Interestingly, dimer–Gs binding exhibited both a slower association rate (*k*_on_) and dissociation rate (*k*_off_) compared to monomer–Gs binding, while AP’s effect on the monomer only reduced *k*_on_ (Table S2). These results suggest that AP partially inhibits recruitment of Gs by β_2_AR, with a more pronounced effect when tightly bound within the dimer. The smaller *k*_off_ observed in dimer–Gs dissociation may reflect a cooperative mechanism, where Gs engagement with one protomer promotes a Gs-favorable conformation in the adjacent protomer, thereby increasing Gs residence time during dissociation.

We further performed radioligand binding assays to assess the allosteric effect of AP on orthosteric ligand binding using isolated monomer or dimer reconstituted in nanodiscs. The AP-bound dimer showed an ∼5-fold higher affinity for [^3^H]-dihydroalprenolol (DHA), a β_2_AR antagonist, in the saturation binding, while AP at 10 μM did not significantly affect [^3^H]-DHA binding to the monomer (Figure S11A). In agonist competition assays with Iso, the dimer exhibited a ∼5-fold increase in the inhibitory constant (*K*_i_), but no significant effect of AP on the monomer (Figure S11B). These findings suggest that AP stabilizes β_2_AR in a more inactive-like conformation in the dimer. We next evaluated the potency of Gs in stabilizing the active state of β_2_AR by allosterically modulating the orthosteric pocket to reduce antagonist binding. We titrated Gs(GDP) into β_2_AR pre-bound with saturated [^3^H]-DHA and quantified the remaining bound [^3^H]-DHA. The results showed that Gs modulates the orthosteric pocket in both dimeric and monomeric β_2_AR, irrespective of AP presence (Figure 5E). This suggests that while AP biases β_2_AR toward an inactive-like conformation, it does not significantly hinder Gs engagement at equilibrium. We then assessed the effects of AP on the function of β_2_AR as a guanine nucleotide exchange factor (GEF), which facilitates GDP-GTP exchange in Gs and accelerates GTP hydrolysis. Using the GTPase-Glo™ assay to monitor GTP turnover, we found that while AP significantly slows down the GTP turnover rate of monomeric β_2_AR, it did not affect the final efficacy. However, we observed much stronger inhibition in GTP turnover for the dimeric β_2_AR (Figure 5F). This likely stems from two factors: (1) the slower association rate *k*_on_ of Gs with dimeric β_2_AR as observed by BLI (Figure S10F) and (2) AP’s biasing of β_2_AR toward an inactive-like state, potentially introducing additional rate-limiting steps in a full productive cycle.

To test if AP impedes β-arrestin core engagement with β_2_AR, we performed competition radioligand binding assays measuring β-arrestin–induced positive allosteric modulation of agonist binding. Using the sortase ligation strategy, we attached the synthetic phosphorylated vasopressin receptor 2 (V_2_R) peptide (V_2_Rpp) to monomeric or dimeric β_2_AR (Figure 5G and S12A), ensuring full tail engagement of β-arrestin, a prerequisite for efficient core engagement. Monomeric β_2_AR in nanodiscs displayed a ∼7-fold increase in Iso affinity in the presence of β-arrestin1 (Figure S12B), consistent with previous reports^30,31^. AP reduced this increase in affinity to ∼2-fold in the monomer and abolished it in the dimer (Figure 5G), suggesting AP partially hinders β-arrestin core interactions on monomers and completely prevents it in dimers. These data align with structural predictions where steric clashes between β-arrestin and the adjacent protomer in the dimer prevent core engagement.

Previously, we showed that DFPQ inhibits GRK-mediated phosphorylation of β_2_AR, a prerequisite for β-arrestin recruitment, in cells^4^. To identify which species is responsible for this inhibition, we measured GRK5 activity towards purified dimeric and monomeric β_2_AR. AP-bound βCAR dimers exhibited near-complete inhibition of GRK5 phosphorylation in both detergent and nanodiscs (Figure 5H and S13A), whereas GRK5 activity toward monomer in nanodiscs remained unaffected by AP (Figure 5H and S13A). The slight reduction in GRK5 activity observed for monomer in detergent at high AP concentrations (Figure 5I) likely reflects the inefficient dimerization of βCAR in detergent (Figure S1A), as we previously observed (Figure 1C). However, the dose-dependent inhibition observed for monomer reconstituted in bicelles (Figure 5I) is a result of AP-induced dimerization in membrane, underscoring the requirement of a continuous lipid bilayer for effective receptor dimerization. The relatively large diameter of typical bicelles (∼40 nm) compared to nanodiscs (∼12 nm) increases the likelihood of incorporation of multiple receptors and provides a fluid membrane environment conducive to dimer formation. AP-dependent inhibition of GRK5 activity was further corroborated by a BRET-based assay monitoring AP effect on direct GRK5 recruitment to the β₂AR (Figure S13B and S13C). These findings are consistent with structural analyses indicating that steric clashes in the dimer hinder GRK engagement, further supporting the role of the dimer in preventing β-arrestin recruitment.

### Negative allosteric effect of AP on **β**_2_AR conformation dynamics

Modulation of protein dynamics often underpins how ligands achieve biased effects in highly dynamic proteins like GPCRs. To provide further insights into the mechanism of AP’s biased NAM effect, we explored how AP tunes the dynamic behaviors of β_2_AR. To this end, we performed smFRET microscopy to monitor conformational changes and double electron–electron resonance (DEER) spectroscopy to measure residue-level distance distributions. Using the established protocols in our lab^24,32^, we placed fluorescent dyes for smFRET or spin labels for DEER near the intracellular ends of TM4 and TM6 on the minimal cysteine construct of β_2_AR. This allowed us to monitor TM6 outward movement, a hallmark of GPCR activation. Increased distances between TM4 and TM6 sites correspond to high-to-low FRET state transitions in smFRET experiments. As shown in Figure 6A and detailed in the Methods, we used a two-affinity-tag purification strategy to produce an AP-mediated β_2_AR “heterodimer”, where only one protomer contained double-cysteine mutants for dye labeling. The recorded traces from the TM4–TM6 sensors of β_2_AR predominantly fall into two FRET states (Figure 6B), representing an active and inactive state. In the absence of an agonist, β_2_AR was primarily in the inactive state; however, monomers without AP exhibited a significantly higher fraction of active species compared to dimers, which showed almost no active population (Figure 6C and 6D). Adding AP to monomers substantially reduced the active state population from 26.5 ± 2.9% to 11.2 ± 3.2% (Figure 6D), whereas adding Iso to dimers failed to increase active state species (Figure 6C). These results indicate that AP stabilizes an inactive conformation of TM6, and Iso alone fails to drive TM6 transition into an active state in the dimer. This is consistent with the observed enhancement of antagonist binding in the dimer (Figure S11A and S11B). Upon adding Gs to the dimer or the monomer with AP, β_2_AR transitioned predominantly to an active state, although dimers retained a higher fraction of inactive species (Figure 6C and 6D), in line with Gs engaging only one protomer of the dimer, while the other may remain in inactive or partially active conformation. This indicates the potential of Gs to overcome the AP-induced stabilization of the inactive conformation, driving receptor activation even in the dimeric form. Washing Gs away from dimers reversed the population to a primarily inactive state, resembling the condition without Gs coupling (Figure 6C). DEER measurements on the β_2_AR (Figure S14), with spin labels reporting on the TM4–TM6 conformation (Figure 6E), revealed that AP causes changes that are consistent to those observed by smFRET. Specifically, AP biases Iso-bound receptors toward a more homogeneous inactive-like conformation (Figure 6F), while having a minimal effect on receptors bound to the super-agonist BI (Figure 6G). The addition of Gs caused a fraction of the fully active conformer to become populated, regardless of AP presence (Figure 6H). In addition to conformational effects, the modulation depth parameter in pulsed dipolar EPR spectroscopy provides a quantitative measure of spin cluster size, with higher values reflecting increased local concentrations of spin labels. Agonist-bound receptor exhibited a reduced modulation depth relative to the Apo receptor, suggesting that receptor activation is associated with decreased dimerization. Addition of AP to agonist-bound receptor resulted in an increase in modulation depth, consistent with enhanced dimerization (Figure S14). Notably, Gs binding largely inhibited the AP-induced increase in modulation depth, indicating that Gs coupling counteracts the dimer-promoting effect of AP. The magnitude of the shifts in modulation depth indicates slight changes in receptor monomer-dimer equilibrium consistent with a predominantly monomeric receptor, as expected for receptor in detergent. However, the clear trend in modulation depth indicates a correlation between the monomer-dimer and inactive-active equilibria. Together, these findings demonstrate that Gs can couple to both dimers and monomers with AP, driving TM6 outward movement. Moreover, the inhibitory effect of AP on the TM6 outward movement of β_2_AR monomer may also contribute to the level of biased signaling observed. This is supported by the observation that β-arrestin inhibition emerges at lower AP concentrations compared to dimerization in cell-based assays. Notably, the effect of AP on β_2_AR dynamics resembles the previous observation in μ-opioid receptor (μOR), where G protein-biased agonists are less efficient than balanced full agonists in stabilizing TM6 outward movement in both smFRET and DEER studies^33^.

**Figure 6.**
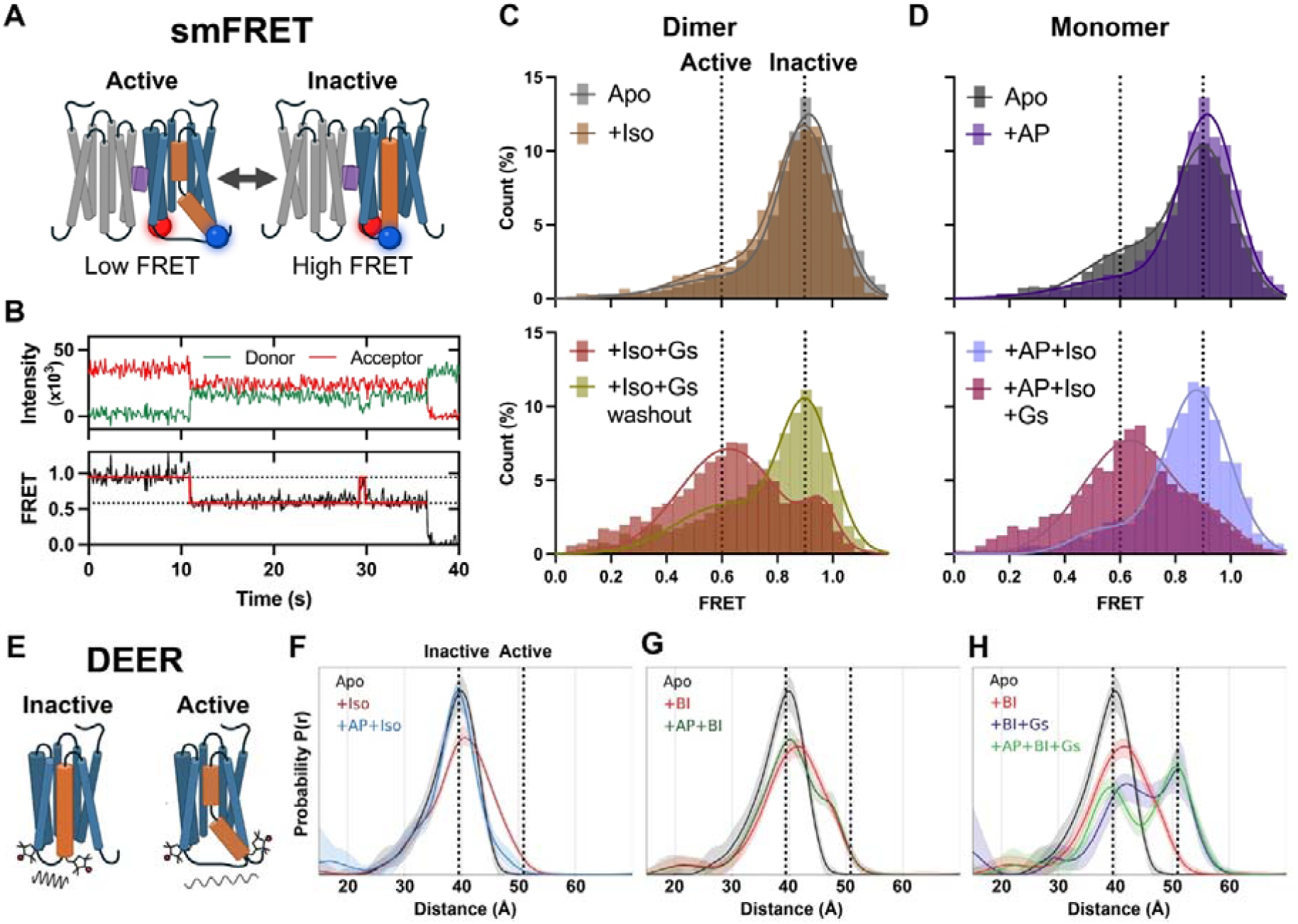
AP biases TM6 conformational dynamics of β_2_AR in both dimer and monomer. (**A**) Schematic of AP-bound β_2_AR dimer labeled on TM4 and TM6 with fluorophores (red and blue spheres) for smFRET analysis. (**B**) Representative fluorescence traces from donor (green) and acceptor (red) channels (top), with corresponding FRET efficiency values (black line) and fitted state model (red line) shown below. Dotted lines indicate distinct FRET states. (**C–D**) FRET population histograms of dimeric (**C**) and monomeric (**D**) β_2_AR showing effects of ligands and Gs on TM6 conformational dynamics. Solid colored lines are two-Gaussian model fits. (**E**) Schematic of β_2_AR with spin labels on TM4 and TM6 for DEER studies. (**F–H**) Distance distributions from DEER measurements of monomeric β_2_AR with spin labels on TM4 and TM6 under different ligand and Gs conditions.

## Discussion

Our cryo-EM structures reveal that AP, a β-arrestin–biased NAM, promotes β_2_AR dimerization, thereby modulating downstream signaling. AP shows strong bias toward β-arrestin inhibition in cell-based assays, and its parent compound, DFPQ, produces sustained relaxation of contracted airway smooth muscle (ASM) in both human ASM cells and mouse airway models by preventing β_2_AR desensitization^4^, suggesting its physiological relevance. We elucidate the mechanistic basis underlying this biased effect: AP robustly induces β_2_AR dimerization. Biochemical and biophysical analyses demonstrate that the dimer acts as a biased signaling species, strongly inhibiting GRK-mediated phosphorylation and β-arrestin recruitment while permitting G protein coupling. Structural analysis indicates that steric hindrance imposed by the adjacent protomer prevents GRK and β-arrestin from adopting their canonical binding modes. These findings offer a novel perspective for achieving signaling bias: besides fine-tuning local conformational states, AP exploits the quaternary structure of receptor to selectively restrict transducer interactions.

Based on our results, we propose an activation model for β_2_AR in the presence of AP (Figure 7). We simplify the energy landscape into two states—an inactive ground state and a fully transducer-bound active state—and outline how AP reshapes this profile. In the absence of AP, the balanced agonist drives the receptor into an active conformation that readily accommodates transducer binding due to a low energy barrier. However, the energy barrier to achieve the fully transducer-bound active state is increased by AP. In the monomer, AP binds a similar pocket as in the dimer, biasing the receptor toward an inactive-like conformation. GRK phosphorylation remains largely intact; however, the reduced allosteric enhancement of agonist binding by β-arrestin suggests a partially destabilized engagement state. AP also slows β_2_AR-catalyzed nucleotide exchange in Gs, indicating a higher activation barrier for Gs, although Gs binding and activation remain largely unaffected, suggesting an unaltered active-state free energy. At higher concentrations, AP promotes stable β_2_AR dimerization, resulting in near-complete loss of GRK phosphorylation and blockade of β-arrestin core engagement. This reflects not only increased energy barriers but also a disrupted β-arrestin-bound state, due to steric clashes with the adjacent protomer. GTP turnover by Gs is further slowed in the dimer, again reflecting a higher activation barrier, while the free energy of the Gs-bound state remains similar to that of the monomer. In cells, where monomers and dimers coexist, AP shifts the receptor ensemble toward the more strongly biased dimeric form in a concentration-dependent manner. Despite reduced GTP turnover by Gs, signal amplification likely compensates and enables near-maximal cAMP production even under conditions where dimer is dominant. Thus, the loss of GRK and β-arrestin engagement in the dimer biases the receptor exclusively toward sustained Gs signaling, highlighting dimerization as a mechanism to selectively rewire family A receptor signaling.

**Figure 7.**
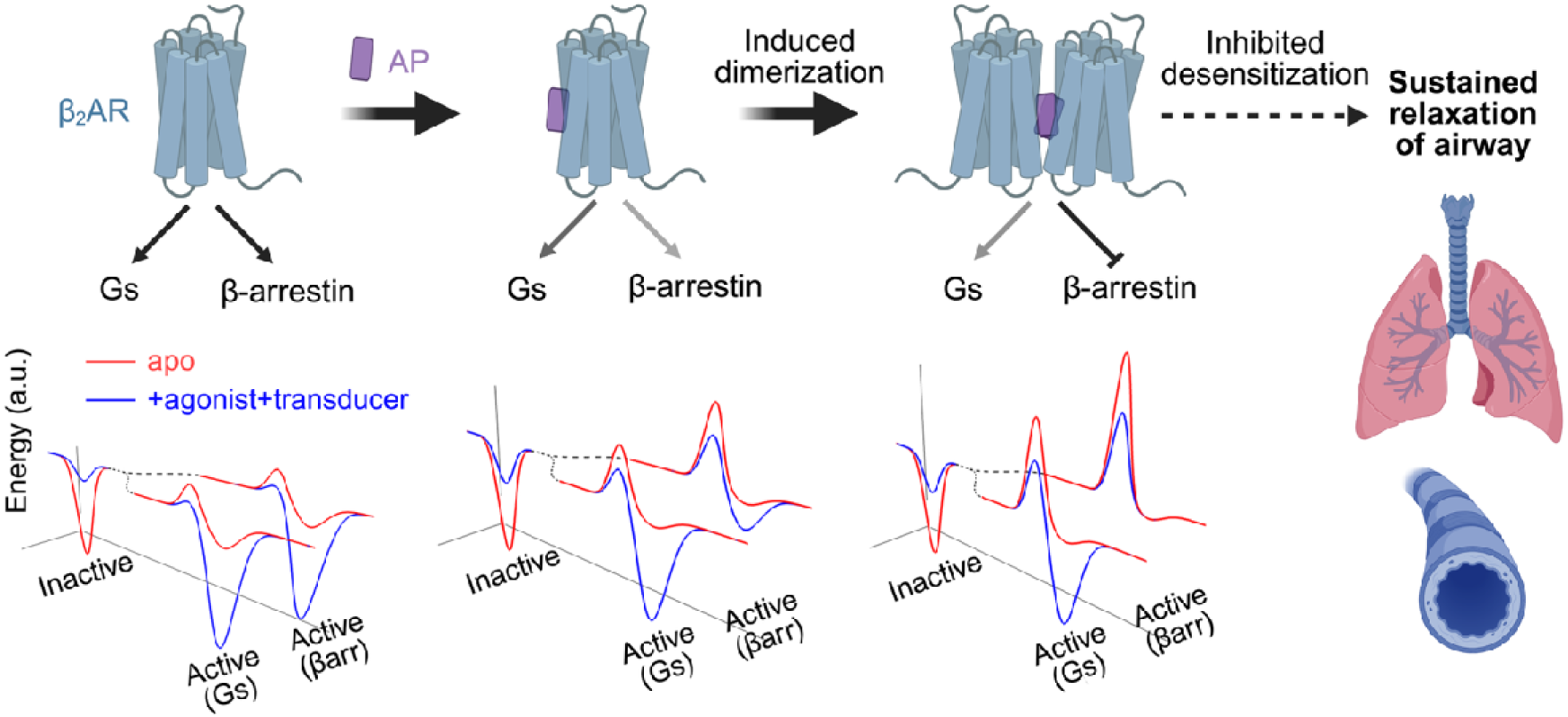
Proposed model for AP biasing β_2_AR signaling. AP (purple) promotes β_2_AR dimerization, which reduces receptor desensitization in human ASM cells and mouse airway models. Simplified free-energy landscapes illustrate how AP alters activation energy barriers and state equilibria of the apo (red) and agonist/transducer-bound (blue) conditions under three scenarios: β_2_AR without AP (left), AP-bound monomer (middle), and AP-bound dimer (right).

The dimer interface observed in our AP-bound β_2_AR complex displays unique features compared to other family A GPCR dimers. While previous studies have reported dimer structures in receptors such as APJR^12,18^ and rhodopsin^1^ by cryo-EM, our AP-stabilized dimer predominantly involves interactions on TM3, TM4, TM5, and ICL2, a configuration distinct from the earlier β_2_AR crystal structure^17^ and other family A GPCR dimers. Importantly, the outward mobility of TM5 and TM6 is not restricted in our dimer configuration, which is essential for efficient transducer engagement. Notably, recent work^34^ on the platelet-activating factor receptor using a cysteine crosslinking strategy reported that dimerization significantly biases signaling toward G protein pathways while limiting β-arrestin recruitment; one of their proposed dimer models resembles our structure with a TM3–TM4–TM5 interface. Similarly, recent studies used a computational and mutagenesis approach to design stable TM4–TM5 dimers of CXCR4, which biased signaling towards Gi^35^.

Beyond discrete dimer formation, cryo-EM imaging reveals that AP-mediated β_2_AR dimerization drives the assembly of larger nanoclusters, and our structural model of the dimer-of-dimer provides a potential molecular basis for specific interactions that underlie this mesoscale organization. Such nanoclusters likely serve as dynamic signaling platforms that modulate both the amplitude and duration of receptor-mediated responses^25^. Recent papers^36–38^ on higher-order transient structures (HOTS) highlight that dynamic oligomerization is a conserved strategy in membrane proteins, which may be beneficial for signaling efficiency and specificity in cells. Consistent with HOTS, AP-induced β_2_AR nanoclusters may fine-tune the balance between receptor activation, desensitization, and internalization. Although our GTP turnover assays on isolated AP-mediated dimers showed reduced turnover activities, the enlarged nanoclusters observed in cells may locally concentrate G proteins, thereby supporting more productive signaling events. Further investigations into the physiological relevance and functional consequences of these nanoclusters will help uncover the potential for harnessing such mechanisms across other receptor systems.

Our findings extend the emerging paradigm of molecular glues in pharmacology^39,40^. AP functions as a GPCR molecular glue by bridging two β_2_AR protomers to form a dimeric assembly with biased functions. Although molecular glues that stabilize GPCR interactions with transducers^41^, including G proteins^42,43^, β-arrestin^44,45^, and GRK^29^, have been reported, AP represents the first example of a ligand mediating TM interactions that drive dimerization of a family A GPCR. This mechanism is reminiscent of that observed in the membrane-bound molecular glue, NVS-STG2^46^, for stimulator of interferon genes (STING). NVS-STG2 induces high-order oligomerization of STING, robustly triggering downstream immune signaling^46^. Both examples illustrate how small molecules can harness and reshape the intrinsic propensity of membrane proteins to form oligomeric assemblies, thereby selectively modulating signal transduction.

In summary, our study deciphers the structural basis by which AP acts as a biased NAM for β_2_AR and provides broader insights into how receptor dimerization impacts GPCR signaling. Such insights not only deepen our understanding of structure–function relationships in GPCR modulation but also highlight potential strategies for designing next-generation ligands that exploit oligomerization as a route to bias receptor signaling.

## Resource availability Lead contact

Further information and requests for resources and reagents should be directed to the lead contact, Brian K. Kobilka.

## Materials availability

Plasmids for minimal cysteine constructs of β_2_AR are available upon request.

## Data availability

The cryo-EM map for β_2_AR_dimer(AP) in nanodisc has been deposited to the Electron Microscopy Data Bank under the accession code <u>EMD-72202</u>. The corresponding atomic model has been deposited in the Protein Data Bank under the accession code <u>9Q3L</u>. Any additional data reported in this paper are available from the lead contact upon request.

## Acknowledgements

We thank R. Qiu of R. S. Lewis’s lab for help with smFRET experiments, E. S. Bruguera for instructions on BLI measurements, M. R. Eckart and J. Tran of Stanford Protein and Nucleic Acid (PAN) Facility for SPR data collection, P. A. N. Reddy and J. M. Salvino for providing AP-7-168, and B. Singal and C. Zhang for support on cryo-EM data collection at the Stanford cryo-EM center (cEMc) and Stanford-SLAC Cryo-EM Center (S2C2), which is supported by the US National Institute of General Medical Sciences (1R24GM154186). We also thank M. Bouvier for providing pcDNA-β-arrestin2-GFP10 and pcDNA3-β_2_AR-RlucII, L. I. Jiang for providing the cAMP intramolecular BRET sensor CAMYEL, G. Milligan for providing pcDNA3-β_1_AR-GFP, and P. S. Chae for providing TTG-T10 detergent. J. Xu is an investigator of SUSTech Institute for Biological Electron Microscopy. This research was supported by National Institutes of Health (NIH) awards R35NS137408 (B.K.K.), R01GM083118 (B.K.K.), P01HL114471 (J.L.B.), R01AI161296 (J.L.B.), R01GM135581 (M.T.L.), S10OD025260 (M.T.L.), and American Heart Association (AHA) Postdoctoral Fellowship 25POST1411512 (J.S.).

## Methods

### Expression and Purification of β_2_AR in Sf9 cells

The β_2_AR construct PN1 was expressed and purified as previously described^32,47^. Briefly, Sf9 cells were infected with a PN1-containing baculovirus produced via BestBac method. Cells were then harvested and resuspended in chilled lysis buffer containing 10 mM HEPES, pH 7.4, 1 mM EDTA, 1 μM alprenolol, and protease inhibitors (leupeptin and benzamidine). Lysed cells were then pelleted at 18,600 rpm for 20 minutes and dounced to homogeneity in chilled solubilization buffer containing 20 mM HEPES pH 7.4, 350 mM NaCl, 1% n-dodecyl β-D-maltoside (DDM), 0.1% cholesteryl hemisuccinate (CHS), 2 mM MgCl_2_, 1 μM alprenolol, protease inhibitors, and benzonase. After stirring for 90 minutes at 4 °C and centrifugation at 18,600 rpm for 30 minutes, 2 mM CaCl_2_ was added to the soluble fraction, which was then applied to anti-FLAG (DYKDDDDK) M1 immunoaffinity resin. The receptor was then washed (20 mM HEPES, pH 7.4, 350 mM NaCl, 0.1% DDM, 0.01% CHS, 2 mM CaCl_2_, leupeptin, and benzamidine), eluted (20 mM HEPES, pH 7.4, 350 mM NaCl, 0.1% DDM, 0.01% CHS, 5 mM EDTA, and 200 ug/mL FLAG peptide), and further purified on Superdex 200 10/300 Increase gel filtration column equilibrated in NH buffer (20 mM, HEPES pH 7.4, 100 mM NaCl) plus 0.1% DDM and 0.01% CHS. To produce homogeneous β_2_AR dimer, Sf9 cells expressing PN1 were resuspended at room temperature (RT) in 20 mM HEPES, pH 7.4, 150 mM NaCl, 10% glycerol, 1 μM alprenolol, protease inhibitors, and 10 μM AP. After a 30-minute incubation at RT, membrane solubilization was initiated by adding 1% lauryl maltose neopentyl glycol (LMNG) and 0.1% CHS. The purification steps followed the same protocol as for the monomeric receptor, except that 0.01% LMNG replaced DDM in all buffers, and 10 μM AP was maintained throughout the purification.

### Expression and Purification of heteromeric G**α**s**β**1**γ**2

As previously described^48,49^, heterotrimeric Gs was expressed and purified from *Trichoplusia ni* Hi5 cells. Briefly, two baculoviruses were generated from the BestBac method, one encoding the wild-type human Gαs subunit and the other encoding the wild-type human β1γ2 subunits containing a histidine tag on the N-terminus of the β subunit. Cells were infected with both viruses for 48 hours and harvested via centrifugation. The pellet was then resuspended and stirred for 30 minutes at 4 °C in hypotonic buffer containing 10 mM HEPES pH 7.4, 100 μM MgCl_2_, 5 mM β-mercaptoethanol (BME), 20 μM GDP, and protease inhibitors. Lysed cells were then pelleted at 18,600 rpm for 15 minutes and dounced to homogeneity in chilled NH buffer plus 1% sodium cholate, 0.05% DDM, 1 mM MgCl_2_, 5 mM BME, 20 μM GDP, and protease inhibitors. After solubilization for 1.5 hours while stirring at 4 °C and centrifugation at 18,600 rpm for 35 minutes, 20 mM of imidazole was added to the soluble fraction, which was then allowed to batch-bind to washed nickel-chelated Sepharose for 2 hours. Pelleted resin was then loaded into a narrow column, washed with buffers containing gradually declining amounts of cholate, and eluted with NH buffer plus 0.05% DDM, 1 mM MgCl_2_, 20 μM GDP, 100 μM tris(2-carboxyethyl)phosphine (TCEP), and 250 mM imidazole. Human rhinovirus 3C protease was added to cleave the histidine tag and the eluate was dialyzed overnight at 4C in 2 L of dialysis buffer (NH buffer plus 1 mM MgCl_2_, 0.05% DDM, 20 μM GDP, and 100 μM TCEP). The protein solution was run through a second nickel-chelated Sepharose column, washed with dialysis buffer supplemented with 20 mM imidazole, and dephosphorylated for 30 minutes on ice with lambda protein phosphatase, calf intestinal phosphatase, and Antarctic phosphatase with 1 mM manganese chloride. The heterotrimer was further purified from excess βγ subunits by using ion exchange chromatography on a MonoQ 10/100 GL column. Sample was loaded and washed with 20 mM HEPES, pH 7.4, 1 mM MgCl_2_, 0.05% DDM, 100 μM TCEP, and 20 μM GDP. Heterotrimer Gs was then eluted with a linear salt gradient from 50 mM NaCl to 500 mM NaCl.

### Expression and Purification of Nb60

Nb60 was expressed and purified as previously described^15^. Briefly, Nb60 was expressed in *Escherichia coli* BL21(DE3) cells. Lysate was then purified on nickel-chelated Sepharose column and subsequently on a Superdex 200 Increase 10/300 column in 20 mM HEPES, pH 7.4, and 150 mM NaCl.

### Sample preparation for cryo-EM

For the sample in detergent, purified PN1 in 0.1% DDM/0.01% CHS was loaded onto anti-FLAG M1 immunoaffinity resin equilibrated in the same detergent mixture supplemented with 2 mM CaCl_2_. A detergent exchange was then performed, during which the receptor bound to the resin was washed with increasing ratios of TTG-T10 to DDM, ultimately transitioning the receptor into 0.01% TTG-T10/0.001% CHS. Each detergent exchange buffer contained NH buffer with 2 mM CaCl_2_, 1 μM carazolol, and 10 μM AP. The receptor was subsequently eluted in NH buffer plus 0.01% TTG-T10, 0.001% CHS, 1 μM carazolol, 10 μM AP, 5 mM EDTA, and 200 ug/mL FLAG peptide. After the detergent exchange, PN1 was incubated for 1 hour with 2-fold molar excess of Nb60. Excess Nb60 was cleared on anti-FLAG M1 immunoaffinity resin after washing with NH buffer plus 0.0006% TTG-T10, 0.00006% CHS, 1 μM carazolol, 10 μM AP, and 2 mM CaCl_2_. The complex was then eluted off the resin with NH buffer plus 0.0006% TTG-T10, 0.00006% CHS, 1 μM carazolol, 10 μM AP, 5 mM EDTA, and 200 μg/mL FLAG peptide. The final sample was concentrated to >10 mg/mL and used immediately for cryo-EM grid preparation.

To prepare the AP-bound β_2_AR dimer in lipid nanodiscs, the purified dimer in LMNG was reconstituted into nanodiscs following a previous protocol^47^ with modifications. Lipids were prepared by mixing 1-palmitoyl-2-oleoyl-glycero-3-phosphocholine (POPC, Avanti), 1-palmitoyl-2-oleoyl-sn-glycero-3-phospho-L-serine (POPS, Avanti), and cholesterol (Sigma-Aldrich) at a molar ratio of 7:2:1, followed by drying under argon and vacuum desiccation for 2 hours. Lipids were resuspended in NH buffer containing 14 mM DDM at 20 mg/mL. The receptor was diluted to 10 μM, incubated with 50 μM BI (MedChemExpress) for 10 minutes on ice, and then combined with membrane scaffold protein (MSP) 1E3D1 and lipids at a molar ratio of 1:2.5:100. The mixture was incubated on ice for 1 hour to allow nanodisc assembly. Detergent was removed by sequential addition of semi-wet Bio-Beads SM2 (60 mg/mL) three times over 3-hour period, followed by overnight incubation at 4°C. The next day, Bio-Beads were removed, and empty nanodiscs were separated by M1 affinity purification. The dimeric βDAR in nanodiscs were eluted in buffer containing 1 μM BI, 10 μM AP, 5 mM EDTA, and 200 μg/mL FLAG peptide. The eluate was further polished by SEC in NH buffer plus 1 μM BI and 10 μM AP. Peak fractions were pooled and concentrated to 5 mg/mL for grid freezing.

The cryo-EM grids were prepared using Vitribot Mark IV (Thermo Fisher Scientific). Quantifoil R1.2/1.3 Au grids were glow-discharged with air for 90 s at 10 mA using Plasma Cleaner (PELCO EasiGlowTM). Aliquots of 3 μL protein sample were applied to the glow-discharged grids. After blotting with filter paper (Ted Pella, Inc) for 3.0 s, the grids were plunged into liquid ethane cooled with liquid nitrogen.

### Cryo-EM data collection and processing

For detergent- and nanodisc-reconstituted β_2_AR samples, a total of 7,139 and 17,997 micrograph stacks, respectively, were collected using a Titan Krios G3i operating at 300 kV on a Falcon 4i direct electron detector or on a K3 camera (Gatan) with a Quantum energy filter. Micrographs were recorded at a nominal magnification of 130,000×, corresponding to a calibrated pixel size of 0.95 Å (detergent) or 0.83 Å (nanodisc) with defocus values ranging from –2.0 μm to –0.8 μm. Each movie stack received a total electron dose of ∼50 eD/Å² over 40 frames.

Motion correction and contrast transfer function (CTF) estimation were performed using the patch motion and patch CTF in cryoSPARC^50^. Particle picking was carried out using the blob picker, and extracted particles were binned 4× for initial 2D classification. 2D classes with recognizable structural features were manually selected, and monomeric and dimeric particles were grouped separately. Ab initio reconstruction was performed using ∼10% of the selected particles to generate four initial 3D references. Iterative rounds of heterogeneous refinement were conducted until poor-quality classes accounted for less than 5% of input particles. For the detergent dataset, 48,805 dimeric and 25,201 monomeric particles were selected for non-uniform (NU) refinement^51^. For the nanodisc dataset, 237,408 particles of dimeric species were selected. Particles were imported into RELION 4^52,53^ for Bayesian polishing^54^ and subsequently returned to cryoSPARC for further NU refinement. C2 symmetry was imposed for the final NU refinement of dimer species After handedness correction, local and CTF refinement were performed using a soft mask centered on the transmembrane domains. Final resolutions were estimated using the gold-standard Fourier shell correlation (FSC) 0.143 criterion. Local resolution estimation was carried out in cryoSPARC.

### Model building and refinement

Initial models of β_2_AR (PDB ID: 2RH1) were docked into cryo-EM maps using UCSF Chimera^55^ and manually adjusted in COOT^56^ to fit the density, including placement of ligands. Real-space refinement was performed in PHENIX^57,58^ with secondary structure and geometry restraints. Model validation was conducted using EMRinger^59^ to assess side chain density fitting. Structural figures were prepared using PyMOL and ChimeraX^60^.

### Mass photometry

Mass photometry measurements were performed using a Refeyn TwoMP instrument (Refeyn Ltd.) and the AcquireMP software (v2.3) following an established protocol^61^. Microscope coverslips (24 × 50 mm, #1.5 thickness; Corning) were cleaned with deionized water and isopropanol, then dried prior to use. Silicone gaskets were applied to the coverslips to form individual wells immediately before sample loading. The instrument was calibrated using NativeMark unstained protein standards (Thermo Fisher Scientific) following the manufacturer’s instructions. Each measurement was conducted by first pipetting 10 μL of NH buffer into a well, followed by focal alignment and locking. Then, 1 μL of β_2_AR sample at a final concentration of 20 nM after dilution was added, mixed gently, and data were acquired for 60 seconds. At least 2,000 binding events were recorded per sample. Data processing and molecular mass determination were performed using the DiscoverMP software suite (Refeyn).

### Cell unroofing and immunogold cryo-EM

Cell unroofing and immunogold labeling were performed as previously described^36,37,62^, with modifications. Quantifoil R1.2/1.3 300-mesh gold EM grids were glow-discharged for 15 seconds, rinsed three times with 70% ethanol, and washed four times with DPBS. Grids were incubated with 0.1 mg/mL poly-D-lysine (Gibco) for 1 hour at RT, followed by four washes in Dulbecco’s phosphate-buffered saline (DPBS, Gibco). Laminin (Sigma-Aldrich, 15 μg/mL) was applied to the grids and incubated at 37°C for 2 hours, then washed again with DPBS. HEK293F cells in suspension were seeded onto prepared grids and cultured until reaching 60–70% confluency. β_2_AR was overexpressed via baculovirus transduction following the BacMam protocol (Thermo Fisher Scientific). Around 16 hours after transfection,cells were rinsed with DPBS containing calcium and magnesium. Grids bearing adherent cells were held with tweezers and dipped into a hypotonic swell buffer (6 mM HEPES-KOH pH 7.4, 43.3 mM K-gluconate, 1.6 mM NaCl, 0.6 mM MgCl_2_) for 30 seconds. An additional 6 μL of swell buffer was added to each grid, followed by blotting with Whatman Grade 5 filter paper (Sigma-Aldrich) to remove the apical membrane. Unroofed samples were blocked with 3% goat serum (Thermo Fisher Scientific) in DPBS containing protease inhibitors (Thermo Fisher Scientific) for 20 minutes at RT. Primary antibody against β_2_AR (Thermo Fisher Scientific) diluted in DPBS with 1% goat serum was applied for 1 hour. After three washes in DPBS, grids were incubated with a goat anti-mouse secondary antibody conjugated to colloidal gold (Ted Pella, Inc) diluted in DPBS with 1% goat serum for 1 hour, then washed again three times.

Grids were plunge-frozen in liquid ethane using a Leica EM GP2 system and stored in liquid nitrogen. Cryo-EM imaging was performed on a Glacios G2 operated at 200 kV, equipped with a Falcon 4i detector, at a nominal magnification of 11,000×.

### Cell-based BRET

To evaluate the effects of AP on cAMP production and β-arrestin recruitment, HEK293 cells endogenously expressing β_2_AR were transiently transfected with either the BRET-based intramolecular cAMP sensor CAMYEL or with plasmids encoding β-arrestin2–GFP10 and β_2_AR–RlucII. CAMYEL comprises both donor and acceptor fused to the cAMP-binding domain of EPAC and undergoes a conformational change upon cAMP binding that alters the BRET signal^63^. Forty-eight hours post-transfection, cells were pre-incubated with increasing concentrations of AP (0.03– 100 μM) for 30 minutes, followed by stimulation with 1 μM isoproterenol for 30 minutes in the presence of 5 μM coelenterazine H (for CAMYEL) or deep blue coelenterazine (DBC) (Cayman Chemical, Ann Arbor, USA) for the β-arrestin assay. For BRET-based β_2_AR dimerization studies, HEK293 cells were co-transfected with 10 ng/well of either β_2_AR–RlucII, β_2_AR_V129L–RlucII, β_2_AR_TM3–RlucII, or β_1_AR– RlucII (BRET donors), along with 20 ng/well of their respective GFP-tagged counterparts (BRET acceptors). Forty-eight hours post-transfection, cells were incubated with increasing concentrations of AP (0.03–100 μM) or Iso (0.1–100 μM) for 1 hour, followed by addition of DBC substrate and incubation for 20 minutes. To control for potential AP autofluorescence effects on the BRET signal, HEK293 cells transfected with β_2_AR–GFP were treated with AP (0.03–100 μM) for 30 minutes, and the induced GFP signal was directly measured and represented as fold change over basal.

For saturation studies of β_2_AR homodimerization, cells were co-transfected with a fixed amount of β_2_AR–RlucII (10 ng/well) and increasing amounts of GFP-tagged β_2_AR plasmid (0–100 ng/well). BRET measurements were taken 48 hours post-transfection after incubation with DBC for 30 minutes.

To evaluate GRK5 recruitment to the β_2_AR, cells were co-transfected with β_2_AR– Rluc and GRK5–GFP. Cells were either stimulated with increasing concentrations of Iso for 30 minutes followed by DBC incubation for 20 minutes or pre-treated with increasing concentrations of AP (0.03–100 μM) for 1 hour followed by 1 μM Iso stimulation for 30 minutes in the presence of the Rluc substrate DBC.

All BRET signals were recorded at 395 nm (donor emission) and 510 or 530 nm (acceptor emission) using an Infinite F500 plate reader (Tecan, Männedorf, Switzerland). Induced BRET changes were calculated by subtracting the basal BRET signal (in the absence of ligand) from the BRET signal measured after stimulation.

For the AP-mediated cAMP and β-arrestin assays (Figure 1B) as well as GRK5 recruitment, results are expressed as percent of the Iso-alone response. For dimerization studies and Iso-induced GRK5 recruitment, data are presented as percentage of the maximal BRET signal (% of MAX). All dose–response curves were fitted using the log(agonist/inhibitor) vs. response (three parameters) function in GraphPad Prism. Data are shown as mean ± S.E.M., n = 3/4.

### SPR analysis on AP binding

SPR measurements were performed using a Biacore T200 system. Monomeric β_2_AR was captured onto a high-affinity streptavidin (SA) sensor chip (Cytiva) via a biotinylated anti-FLAG M2 antibody (Sigma-Aldrich), yielding a final response of ∼1500 resonance units (RU). The running buffer consisted of 20 mM HEPES (pH 7.5), 100 mM NaCl, 0.01% LMNG. AP was injected at increasing concentrations ranging from 0.6 to 10 μM at a flow rate of 30 μL/min. Association and dissociation phases were recorded for each injection. All sensorgrams were processed using double-referencing to correct for nonspecific binding and instrument artifacts. This was accomplished by subtracting both (i) the response obtained from a blank injection of running buffer over the active surface (to account for bulk refractive index changes and injection artifacts), and (ii) the response from the compound injection over a reference surface without immobilized protein (to correct for nonspecific binding to the surface and matrix effects). Sensorgrams were processed using Biacore Evaluation Software, and binding curves were fit globally using a steady-state affinity model.

### BLI on Gs binding

BLI measurements were conducted at 30°C with continuous shaking at 1000 rpm using an Octet RED384 system (FortéBio). SA biosensor tips (Sartorius) were coated with 10 nM biotinylated anti-FLAG M1 fragment antigen-binding region (Fab) in the NH buffer with 0.01% LMNG and 0.001% CHS for 300 seconds. FLAG-tagged β_2_AR was then captured by incubating the tips in 100 nM β_2_AR with 10 μM Iso for 600 seconds. Following receptor immobilization, the biosensors were transferred into wells containing a concentration series of Gs (30 nM to 10 μM) in the binding buffer containing 10 μM GDP and 0.1% bovine serum albumin (BSA, Sigma-Aldrich) for 180 seconds (association phase), followed by transfer into buffer-only wells for 300 seconds (dissociation phase). Control channels lacking either Gs or immobilized β_2_AR were used for double-reference subtraction. Association and dissociation kinetics were fitted with a single-exponential model to derive apparent *k*_on_ and *k*_off_. Equilibrium binding responses were used to determine the dissociation constant *K*_D_.

### Radioligand binding assay

For saturation binding studies, 50-100 femtomoles of monomeric or dimeric β_2_AR reconstituted in nanodiscs, following the protocol described in the sample preparation for cryo-EM, were incubated with increasing concentration of [^3^H]-DHA at RT for 1 hour in a buffer containing 20 mM HEPES, 100 mM NaCl, 0.5% BSA. Non-specific binding of the radioligand was determined by adding 10 μM alprenolol in the same reaction system. For monomeric β_2_AR, the assay was performed with or without AP. For competition binding studies, monomeric or dimeric β_2_AR reconstituted in nanodiscs were incubated with 1 nM [^3^H]-DHA and increasing concentrations of Iso or GDP-bound Gs in the same buffer as saturation binding. Nanodiscs were separated from excess [^3^H]-DHA on Whatman GF/B filters using a Brandel 96-well harvester. The bound radioligand were read on a liquid scintillation counter (MicroBeta Jet, PerkinElmer). Data were analyzed by GraphPad Prism 10.

### GTP Turnover

The GTPase GLO assay was performed using a modified GTPase-GLO^TM^ assay from Promega as previously described^24,32^. Briefly, 100 nM of monomeric PN1, reconstituted in nanodiscs with MSP1E3D1 following the protocol described in the sample preparation for cryo-EM, was incubated for 1 hour at RT with 20 μM Iso (Sigma-Aldrich) and a range of concentrations of AP in NH buffer plus 0.2% DMSO, and 20 μM GTP. Simultaneously, a 1 μM stock of heterotrimeric Gs protein was prepared in a buffer consisting of NH buffer plus 0.04% DDM, 200 μM TCEP, 20 mM MgCl_2_, and 20 μM GDP. Equal volumes of PN1 and Gs were then mixed and incubated for 60 minutes. The final reaction consisted of 50 nM of ligand-bound PN1 and 500 nM Gs in NH buffer plus 0.1% DMSO, 0.02% DDM, 100 μM TCEP, 10 μM MgCl_2_, 10 μM GTP, and 10 μM GDP. An equal volume of GTPase-Glo reagent in NH buffer plus 0.02% DDM and 5 μM ATP was then added and incubated for 30 minutes. Detection reagent was subsequently added and incubated for 10 minutes. Luminescence was detected using the MicroBeta counter. For the time course assay, the experimental setup remained identical except 200 nM of AP-mediated dimeric PN1 was included as a condition and the PN1-Gs reactions occurred for 30, 60, 90, and 120 minutes.

### Ligation of V_2_Rpp to receptor and β-arrestin competition radioligand binding

β_2_AR constructs modified with a C-terminal sortase recognition sequence (LPETGHH inserted after residue 365) were expressed in Sf9 cells and purified as described above for monomeric and AP-induced dimeric receptors. Sortase-mediated ligation of synthetic V_2_Rpp to receptor was performed as previously described^30,31^. For ligation reaction, 10 μM purified receptor was incubated in NH buffer supplemented with 0.01% LMNG, 0.001% CHS, and 5DmM CaCl_2_ with 50 μM synthetic GGG–V_2_Rpp peptide and 2 μM evolved sortase A pentamutant (eSrtA)^64^. The mixture was incubated overnight at 4 °C. Unreacted receptor and eSrtA (bearing the C-terminal His tag) was removed by binding to nickel-chelated Sepharose resins. Labeled monomeric or dimeric β_2_AR–V_2_Rpp samples were reconstituted into nanodiscs following the protocol described in the sample preparation for cryo-EM.

The equilibrium competition radioligand binding assays were performed with β_2_AR– V_2_Rpp in nanodiscs in the presence of 2DnM [^3^H]-DHA, increasing concentrations of Iso, and 1 μM C-tail-truncated β-arrestin1(382), prepared as previously described^65^. 10 μM AP was added where applicable. After incubation at RT for 1 hour, samples were harvested, and radioactivity was measured as described in the previous section to calculate the *K*i values of Iso.

### GRK5 radiometric phosphorylation assays

To evaluate the effect of β_2_AR dimerization on receptor phosphorylation, β_2_AR monomers or dimers (1 μM), purified in LMNG micelles or reconstituted in nanodiscs following the protocol described in the sample preparation for cryo-EM, were incubated for 5 min at 30°C with purified C-terminally Strep-tagged GRK5 (50 nM) in a reaction buffer containing 20 mM Tris-HCl, pH 7.4, 5 mM MgCl_2_, 30 mM NaCl, 0.5 mM EDTA, 100 μM [γ^32^P]ATP (1,000 to 2,000 cpm/pmol), and 25 μM BI. The β_2_AR samples in LMNG micelles were additionally supplemented with 20 μM C8-PIP_2_ to increase efficiency of β_2_AR phosphorylation in detergent. To evaluate the effect of AP on β_2_AR phosphorylation, purified β_2_AR monomers (1 μM) in LMNG micelles were reconstituted into bicelles with PIP_2_^66^ and AP concentration was varied from 0 to 24 μM. Reactions were quenched with SDS sample buffer, and samples were separated by SDS-PAGE. Gels were stained with Coomassie blue (Sigma-Aldrich), dried, exposed to autoradiography film, and ^32^P-labeled proteins were excised and counted to determine the amount of phosphate transferred. Reaction rates were normalized to phosphorylation of the β_2_AR monomers (β_2_AR monomers and dimers) or to phosphorylation in the absence of AP (AP effect).

### Sample preparation for fluorescence measurements

Site-specific fluorophore labeling of β_2_AR was performed using engineered cysteine mutants on a minimal cysteine background (Δ6), as previously described^24,32^. For smFRET experiments, β_2_ARΔ6 constructs were cloned into the pcDNA-Zeo-tetO vector and transfected into Expi293 cells stably expressing the tetracycline repressor (Thermo Fisher Scientific, A14635). Transfections were carried out using the Expifectamine kit according to the manufacturer’s protocol. Two days post-transfection, receptor expression was induced with 4 μg/mL doxycycline and 5 mM sodium butyrate in the presence of 1 μM alprenolol. Cells were harvested 40 hours after induction and immediately processed for purification.

For studies on the β_2_AR dimer in liposomes, single-cysteine mutants were introduced at TM5 (R228C) or H8 (I334C). Homogeneous dimers were expressed in Sf9 cells and purified as described above. Labeling was performed by incubating 10 μM purified receptor with a 5-fold molar excess of a pre-mixed maleimide-conjugated dye pair: DY549P1 (Dyomics) and Alexa Fluor 647 (Thermo Fisher Scientific) at a 1:1.5 ratio. The reaction was incubated for 30 minutes at RT and quenched with 5 mM L-cysteine. Excess dye was removed by SEC (Superdex 200 Increase 10/300) in 20 mM HEPES (pH 7.4), 150 mM NaCl, and 0.01% LMNG/0.001% CHS. Labeled dimers were reconstituted into liposomes consisting of POPC, 1-palmitoyl-2-oleoyl-sn-glycero-3-phosphoethanolamine (POPE, Avanti) , and cholesterol at a molar ratio of 6:3:1 using an established protocol^67^.

For studies on TM6 dynamics, double-cysteine mutants (N148C on TM4 and L266C on TM6) were introduced. Monomeric receptor was expressed in Expi293 cells and purified following the Sf9 purification procedure. For the dimer, Expi293 cells were co-transfected at a 1:1 plasmid ratio with constructs encoding an 8×His-tagged β_2_ARΔ6 (lacking a FLAG tag) and a FLAG-tagged β_2_ARΔ6 carrying the N148C/L266C mutations. Heterodimers were isolated via tandem affinity purification. Clarified lysates were first incubated with nickel-chelated Sepharose resins and washed with buffer containing 20 mM imidazole. Proteins were eluted with 250 mM imidazole, then supplemented with 2 mM CaCl_2_ and subjected to anti-FLAG M1 immunoaffinity purification. This two-step procedure enriched for heterodimers containing only one protomer with the double-cysteine mutations. Fluorophore labeling was performed as described above for the single-cysteine mutant dimer sample.

### Ensemble fluorescence measurements

Ensemble FRET experiments were conducted in Fluoromax 4C spectrofluorometer (Horiba Scientific) with excitation and emission slit widths set to 5 nm and 3 nm, respectively. Emission spectra were recorded upon excitation at 532 nm. AP-bound β_2_AR dimers labeled with donor and acceptor fluorophores (I334C H8 sensor) were diluted 1,000-fold in NH buffer plus 0.01% LMNG without AP to a final concentration of 1 nM. Fluorescence spectra were collected at 1, 5, 30, 120 minutes, and 16- and 24-hours post-dilution. All spectra were normalized to donor intensity. To assess the effects of transducer binding, samples were incubated with 100 μM Iso and i) 10 μM Gs (in the presence of apyrase) or ii) 20 μM β-arrestin1(382), together with V_2_Rpp and Fab30, which stabilizes the active V_2_Rpp-bound β-arrestin1 conformation. β-arrestin1(382) and Fab30 were prepared as previously described^65^. Samples were incubated for 1 hour in the dark to allow full equilibration prior to measurement. All experiments were performed in triplicate.

### smFRET microscopy

Flow chambers for smFRET experiments were assembled using mPEG-passivated glass coverslips (VWR), doped with biotin-PEG16 (Laysan Bio), as described previously^24,32^. Prior to use, coverslips were incubated with 1 mg/mL NeutrAvidin (Thermo Fisher Scientific), followed by 10 nM biotinylated anti-FLAG M1 Fab. Labeled β_2_AR samples were diluted to 100–500 pM in NH buffer plus 2 mM CaCl_2_ and added to the chambers. After achieving optimal surface density, unbound receptor was washed out using imaging buffer supplemented with 100 μM cyclooctatetraene (Sigma-Aldrich) and an oxygen scavenging system (1% D-glucose, 1 mg/mL glucose oxidase, 0.04 mg/mL catalase).

Fluorescence imaging was performed on a custom-built, objective-based TIRF microscope as reported previously^68^. The setup is built on a Zeiss Axiovert S100 TV platform with a 100×, 1.45 NA oil-immersion objective (Zeiss). Donor and acceptor fluorophores were excited with 532 nm and 637 nm lasers (OBIS LS 150 mW and LX 140 mW, Coherent). Emissions were separated by a 652 nm dichroic beamsplitter (Semrock), filtered through 580/60 nm and 731/137 nm bandpass filters, and split using an OptoSplit II beamsplitter (Cairn Research) onto an EMCCD camera (iXon DU897E, Andor). Data acquisition was controlled by μManager via custom BeanShell scripts, and movies were recorded as stacked TIFFs in frame-transfer mode at 100 ms exposure per frame. Laser power was tuned to balance high signal-to-noise with photobleaching timescales of tens of seconds. Each slide typically yielded 10–20 movies per channel. All imaging was performed at RT.

Fluorescence traces were analyzed using custom Python scripts. Donor and acceptor channels were aligned using registration images, and individual molecules were identified as local intensity maxima within a 5-pixel neighborhood. Donor-only spots were excluded. For each fluorophore pair, intensities were background-corrected using a local circular region (35-pixel diameter). Donor leakage into the acceptor channel (∼7%) was subtracted.

Donor excitation was used to monitor emission for 80 seconds, followed by direct acceptor excitation for 1 second to confirm fluorophore identity. Traces were selected for analysis based on the following criteria: (1) signal-to-noise ratio ≥5; (2) single-step acceptor photobleaching prior to donor bleaching; (3) γ factor between

### 0.5 and 2.5; (4) anticorrelated donor and acceptor intensity fluctuations; and (5) single-step donor bleaching, if present

FRET efficiency (*E*) was calculated as *E = I_a_/(I_a_+*γ*I_d_)*, where *I_a_*and *I_d_* are the background-corrected acceptor and donor intensities, respectively. γ-correction was applied as described previously. For each trace, FRET values were binned into 30 intervals across the range [–0.25, 1.25] and normalized by the total number of data points. Ensemble FRET histograms were generated by averaging the normalized histograms from individual molecules and fit to a two-Gaussian distribution model.

### Sample preparation for DEER

For DEER measurements, β_2_ARΔ6-N148C/L266C was expressed and purified as described above in Sf9 cells. To exchange detergent from 0.1% DDM/0.01% CHS to 0.01% (w/v) LMNG/0.001% CHS, the receptor was extensively washed with a progressive gradient of DDM: LMNG buffer. In parallel, while the receptor was bound to the resin, alprenolol was removed by washing with saturating concentrations of the low-affinity antagonist atenolol. Because of the fast dissociation kinetics of atenolol from the β_2_AR, subsequent washes with ligand-free buffer yielded unliganded β_2_AR for spin labeling. The flag eluted receptor was labeled with the spin label reagent IAP in the presence of 100 μM TCEP in buffer containing 20 mM HEPES, pH 7.4, 150 mM NaCl, and 0.01% LMNG/0.001% CHS. Twenty-fold molar excess of 3-(2-iodoacetamido)-proxyl (IAP) was added to 10 μM β_2_ARΔ6 receptor for 3 hours at RT. After quenching of the reaction with 5 mM final L-cysteine, the receptor was separated from the excess spin label by SEC (Superdex 200 10/300) in SEC buffer (20 mM HEPES, pH 7.4, 150 mM NaCl, and 0.01% LMNG/0.001% CHS) prepared with D_2_O. The sample was concentrated using a 50 kDa concentrator to a concentration > 25 μM. D8-glycerol was added as a cryoprotectant to 25 % (v/v). 13 μL of sample was added to a borosilicate capillary 1.4 mm ID × 1.7 mm OD (VitroCom, Inc) and flash-frozen in liquid nitrogen.

### DEER spectroscopy

DEER experiments were conducted as previously described^32^ at Q-band (∼33.68 GHz) using a Bruker Elexsys 580 spectrometer equipped with a SpinJet AWG, EN5107D2 resonator, variable-temperature cryogen-free cooling system (ColdEdge Technologies Inc.), and a 300 W TWT amplifier (Applied Systems Engineering Inc.). All measurements were performed at 50 K. Dipolar evolution data were acquired using a dead-time-free 4-pulse DEER sequence with gaussian pulses^69^ and with 16-step phase cycling.

The experimental parameters used for DEER data collection were: π/2, π_obs_, and π_pump_ pulse lengths of 40 ns; a frequency offset (Δv) of 90 MHz; d1 = 250 ns; d2 = 5150 ns; shot repetition time = 2000 μs; shots per point = 4; and integration window = 40 ns. The optimal microwave power (i.e., pulse amplitude) for the π/2, π_obs_, and π_pump_ pulses were determined using transient nutation experiments, where pulse amplitudes were adjusted to maximize the inversion of the Hahn echo^70^. Pump pulses were applied to the maximum intensity of the field swept echo detected absorption spectrum. Observe pulses were applied at a frequency 90 MHz lower than the pump pulses.

DEER data were processed using DeerAnalysis 2022^71^, which employs two fitting routines: neural network analysis (DEERNet^72^, Spinach revision 5662) and Tikhonov regularization (DeerLab 0.9.1)^73^. The consensus fit represents the mean of both methods, with reported 95% confidence intervals also incorporating errors from both methods. Time traces were normalized to signal intensity at t = 0, and distance distributions were area normalized. Custom Python scripts were used for plotting the dipolar evolution time traces and the distance distributions.

## Author Contributions

Conceptualization, J.S., T.N.P., J.X., J.L.B., and B.K.K.; methodology, J.S., T.N.P., J.X., K.E.K., F.D.P., and A.M.G.; investigation: J.S., T.N.P., J.X., K.E.K, F.D.P, A.M.G., and H.W.; data curation and formal analysis, J.S., T.N.P., J.X., K.E.K., F.D.P., and A.M.G.; writing – original draft: J.S., J.X., and, T.N.P.; writing – review & editing, all authors; supervision and funding acquisition, B.K.K., J.L.B., and M.T.L.

