## Supplemental information for "A biased allosteric modulator functions as a molecular glue to induce β_2_AR dimerization"

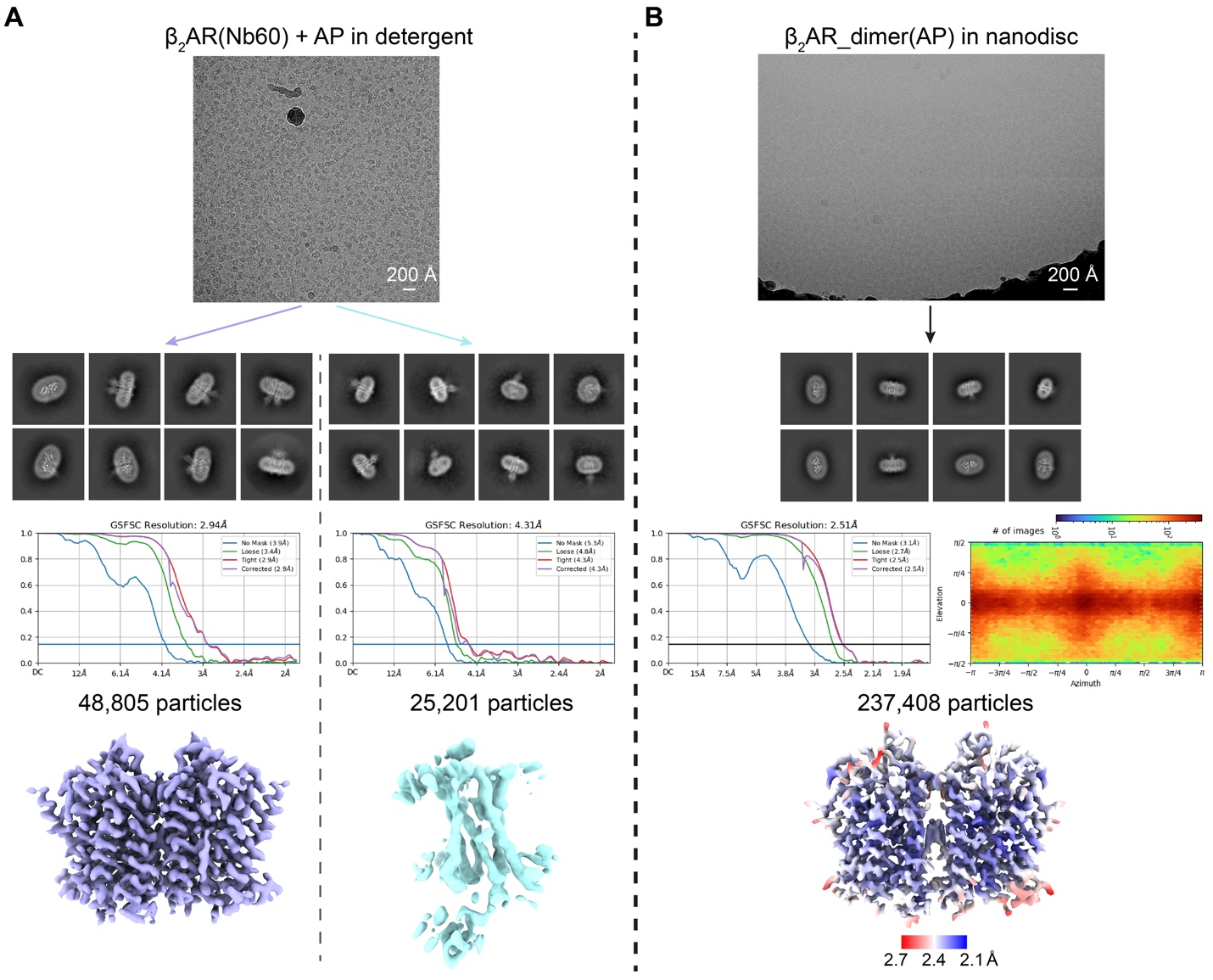

**Figure S1. Cryo-EM structural determination of AP-bound β_2_AR.** Representative micrographs, 2D class averages, gold-standard Fourier shell correlation (GSFSC) curves, and final maps of (**A**) β_2_AR bound to Nb60 in detergent with AP added post-purification and (**B**) AP-bound β_2_AR dimer in nanodiscs. Dimeric and monomeric particles were processed separately after 2D classification to reconstruct final maps of the dimer (lavender) and monomer (cyan) in (**A**). The direction distribution of particles and local resolution heat map for the final reconstruction of β_2_AR(AP)_dimer in nanodiscs are shown in (**B**).

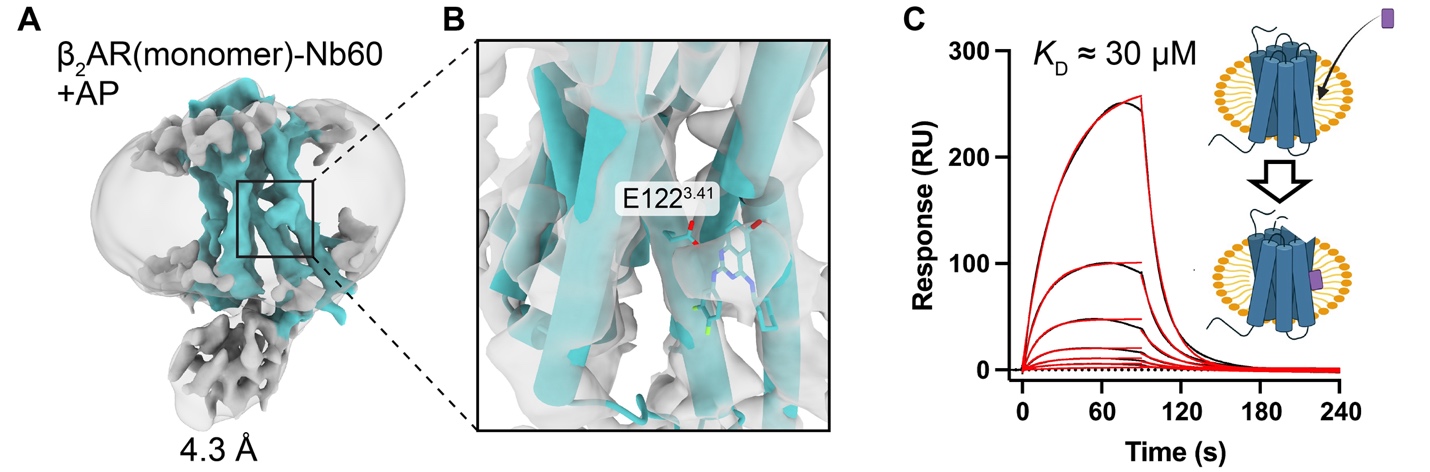

**Figure S2.** **Cryo-EM structure and binding analysis of AP to monomeric β_2_AR.** (**A**) Cryo-EM map of Nb60-bound monomeric β_2_AR in the presence of AP. (**B**) Evidence of AP binding to the monomer at a similar site as in the dimer. One protomer of the dimer (cyan) is fitted into the map. (**C**) SPR sensorgrams (black) with corresponding kinetic fits (red) showing AP binding to surface-immobilized monomeric β_2_AR in detergent. *k*_on_ = 1.92 ×10^3^ M^-1^ s^-1^, *k*_off_  = 6.02 ×10^-2^ s^-1^.

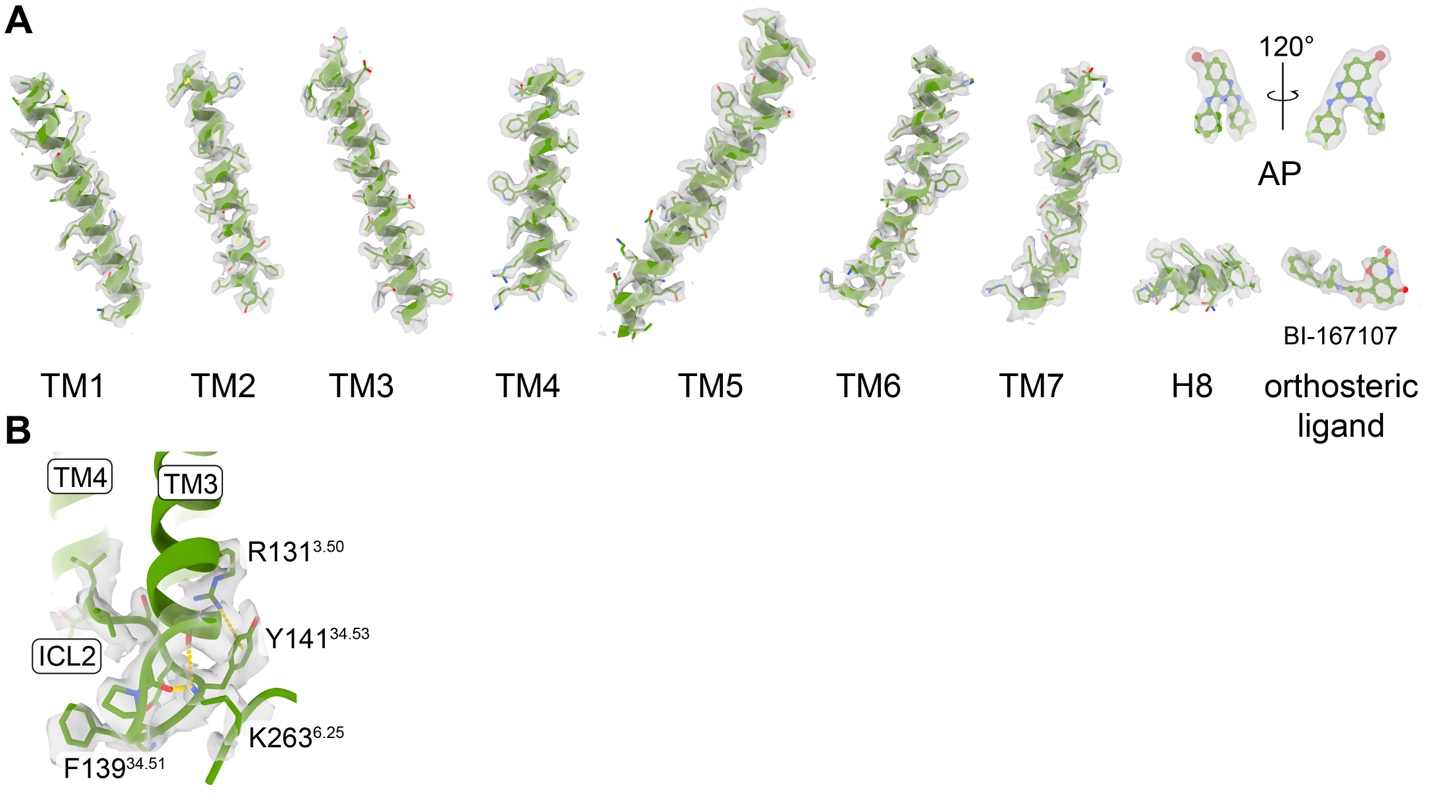

**Figure S3.** **Model-to-map fit of the AP-bound β_2_AR dimer structure in nanodiscs.** (**A**) Side chains of TM1–TM7, H8, AP, and the orthosteric ligand BI from one protomer are shown as sticks with corresponding cryo-EM densities overlaid. (**B**) Conformational rearrangement near ICL2 with EM densities overlaid.

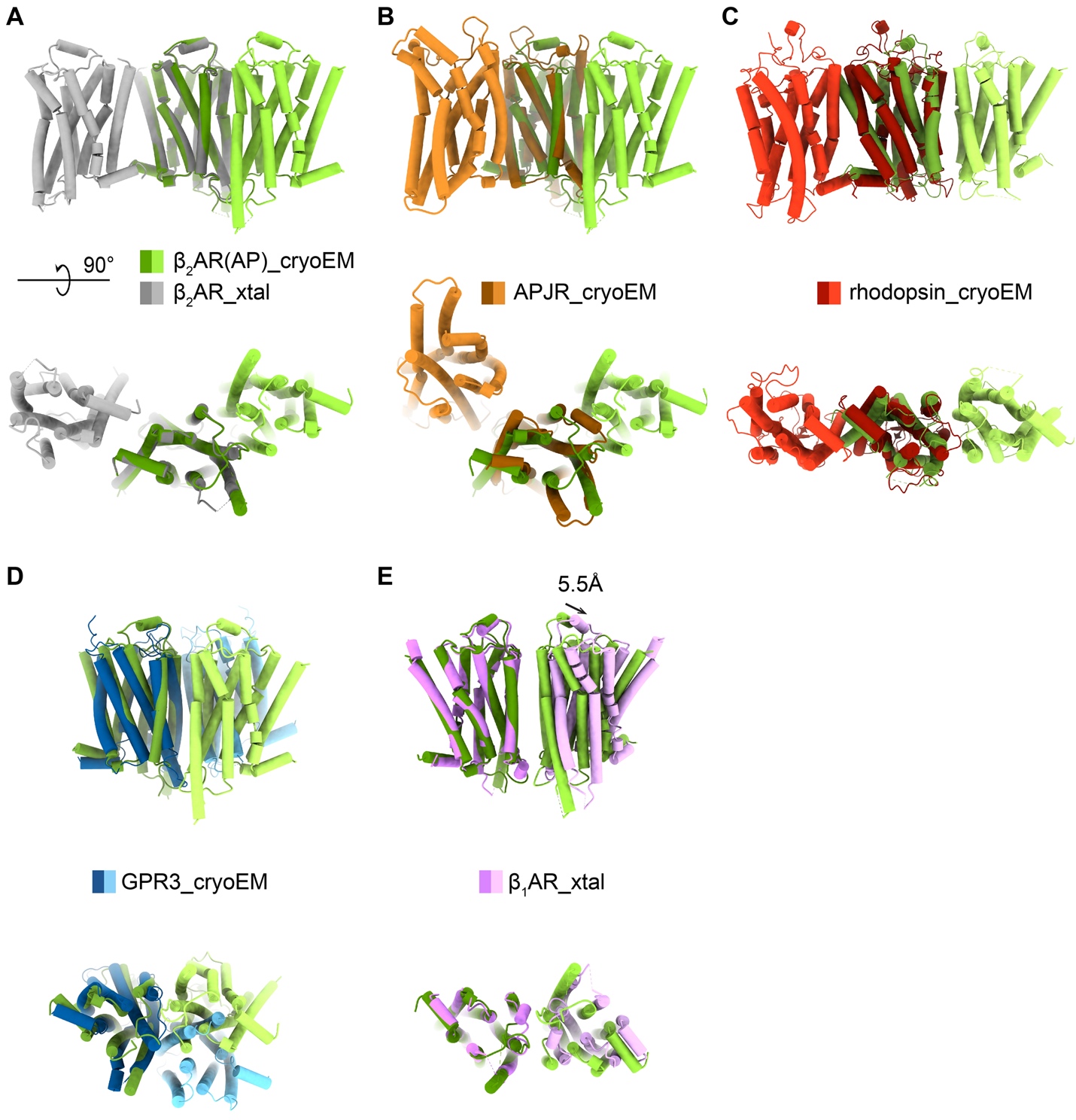

**Figure S4.** **Distinct structural arrangement of AP-bound β_2_AR dimer compared to other class A GPCR dimers.** Structural comparisons with (**A**) the potential dimer observed in the LCP crystal structure of β_2_AR (PDB ID: 2RH1), (**B**) the cryo-EM structure of the human apelin receptor (APJR) dimer (PDB ID: 7W0L), (**C**) the cryo-EM structure of bovine rhodopsin dimer (PDB ID: 6OFJ), (**D**) the cryo-EM structure of human GPR3 (PDB ID: 9LYD), and (**E**) the potential dimer interface of the turkey β_1_AR LCP crystal structure (PDB ID: 4GPO). One protomer from each dimer is aligned for structural comparison. The AP-bound dimer shares similarity with the potential β_1_AR dimer; however, the two protomers in the β_1_AR dimer are positioned ~5.5 Å further apart.

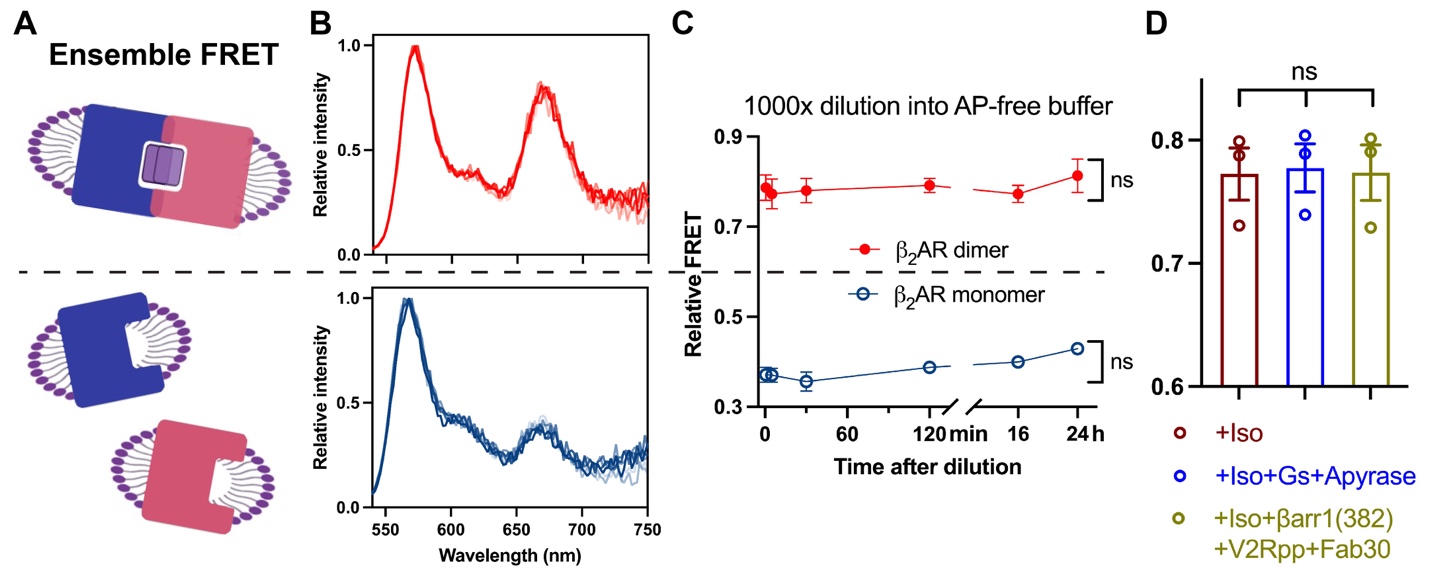

**Figure S5.** **Stability of the AP-bound β_2_AR dimer in solution.** (**A**) Schematic of ensemble FRET assay for receptor dimerization. Red and blue rectangles with an indentation represent receptors labeled with donor or acceptor fluorophores. Purple rectangles represent AP molecules. (**B**) Normalized fluorescence spectra of AP-bound β_2_AR dimer (top) and β_2_AR monomer (bottom) in AP-free buffer. (**C**) Time course of relative FRET for AP-bound β_2_AR dimer (red) and β_2_AR monomer (blue) in AP-free buffer. (**D**) Relative FRET in the AP-bound β_2_AR dimer sample is not affected by agonist, Gs, or activated β-arrestin1. Data are presented as mean with SEM, n = 3. ns, not significant.

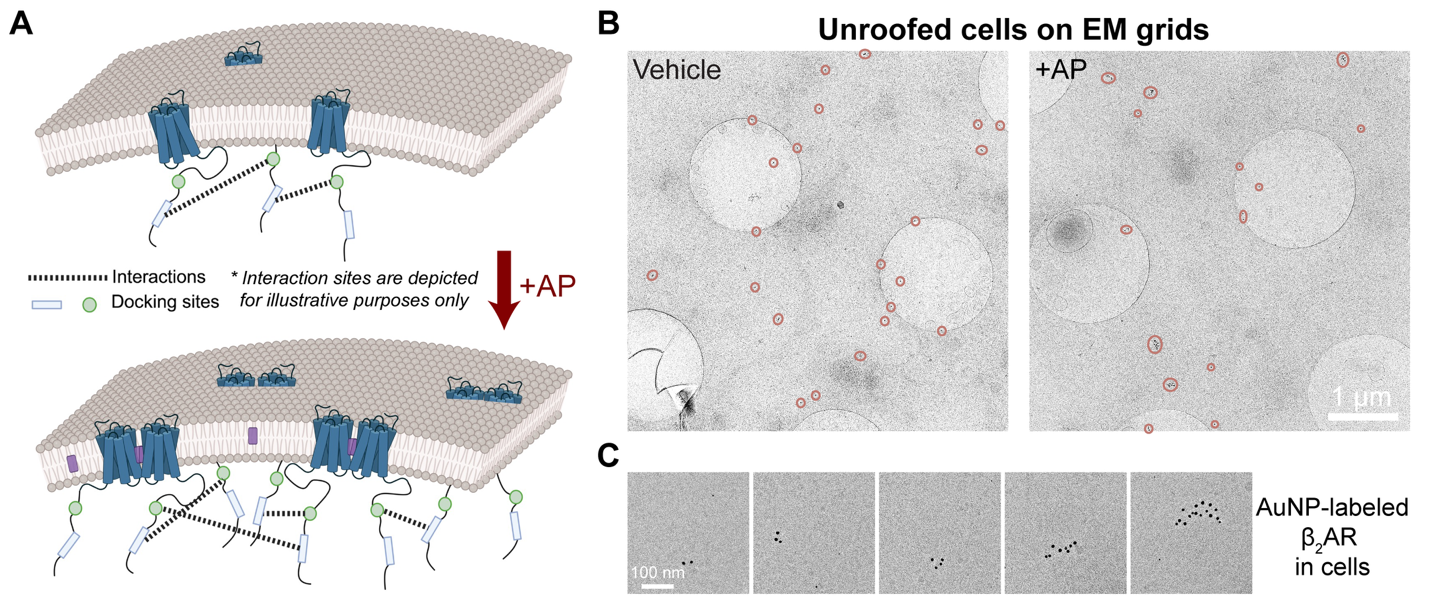

**Figure S6.** **β_2_AR nanoclustering on cell membranes visualized by immuno-gold labeling cryo-EM.** (**A**) Schematic illustrating a potential mechanism by which AP-induced β_2_AR dimerization leads to enlarged nanoclusters. The locations of transient receptor–receptor interactions are shown for conceptual purposes; molecular details remain undefined. Potential contacts observed in **Figure S8C** may contribute to the formation of HOTS. (**B**) Representative cryo-EM images of unroofed cell membranes on grids (Quantifoil R1.2/1.3), showing β_2_AR labeled with gold nanoparticle (AuNP)–conjugated antibodies. Nanoclusters are highlighted with orange circles. (**C**) Gallery of AuNP-labeled β_2_AR nanoclusters displaying varying numbers of particles.

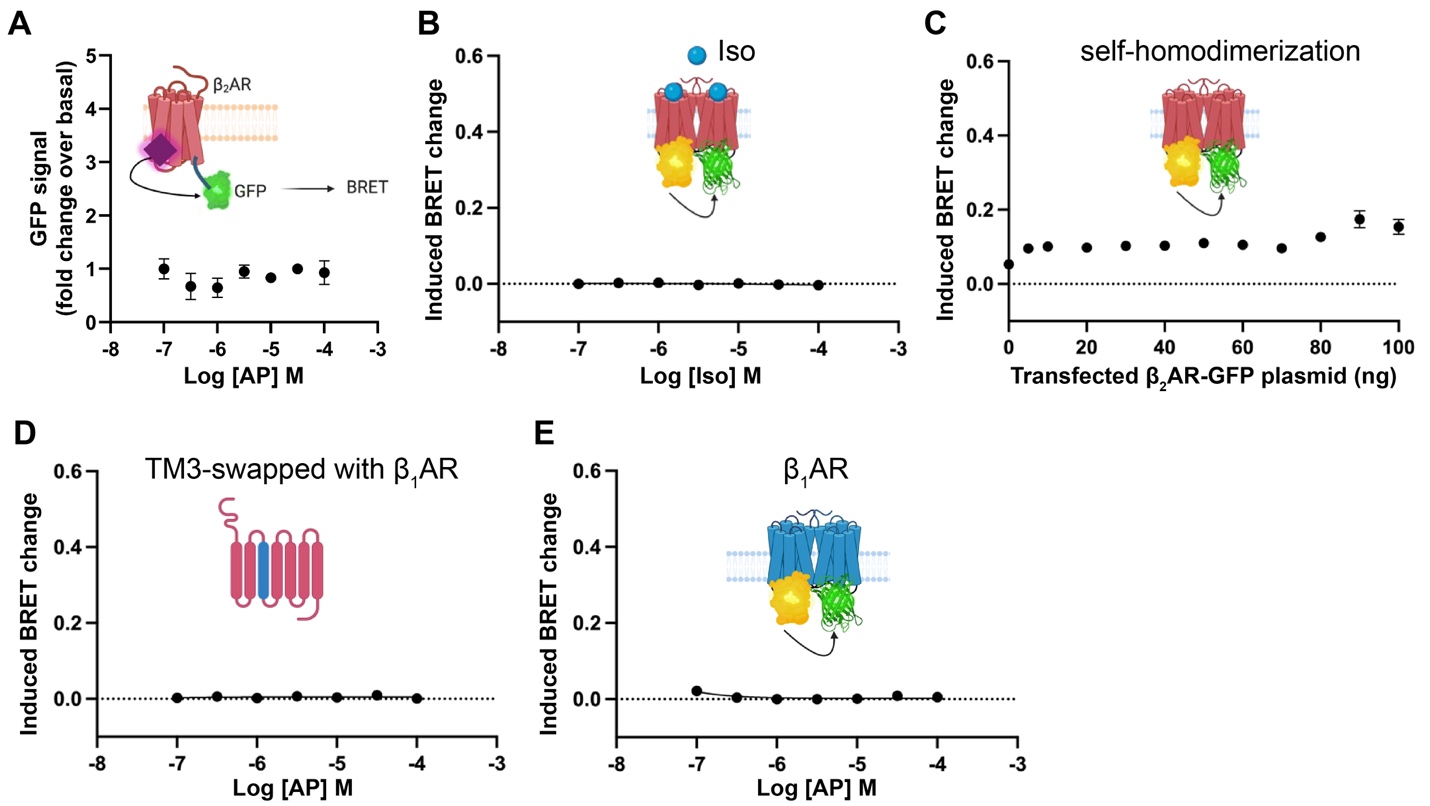

**Figure S7.** **Validation of cell-based BRET assays on receptor dimerization.** (**A**) AP does not generate nonspecific BRET signals. (**B**) Agonist Iso does not induce β_2_AR dimerization in the assay. (**C**) Saturation BRET assays show no significant change in baseline BRET signal with increasing amounts of transfected β_2_AR–GFP plasmid. (**D**) Swapping TM3 of β_2_AR with that of β_1_AR disrupts AP-induced dimerization. (**E**) AP does not induce dimerization of β_1_AR. All data are presented as mean with SEM, n = 3.

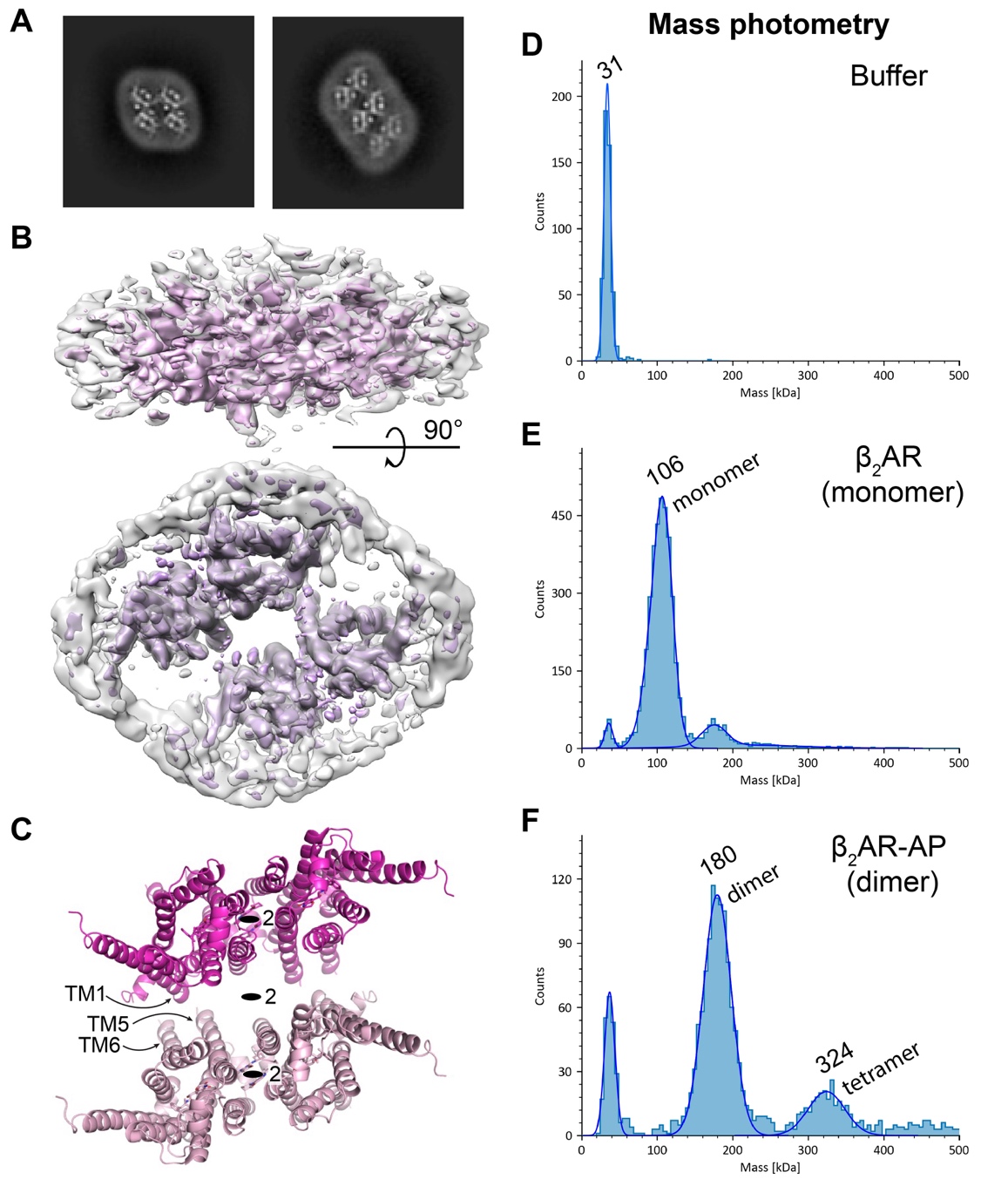

**Figure S8.** **Higher-order oligomers of AP-bound β_2_AR.** (**A**) 2D class averages from the cryo-EM dataset of AP-bound β_2_AR show top views of dimer-of-dimer and trimer-of-dimer species. (**B**) 3D reconstruction of the dimer-of-dimer species. Resolution anisotropy in the map is due to the scarcity of side views. (**C**) Structural model of the dimer-of-dimer, with two β_2_AR dimers docked into the cryo-EM map. TM1–ICL1 and the cytosolic ends of TM5–TM6 form potential inter-dimer contacts. Two-fold symmetry axes are indicated. (**D–F**) Particle mass distributions from mass photometry of (**D**) buffer alone, (**E**) monomeric β_2_AR, and (**F**) AP-bound β_2_AR dimer samples. Samples were diluted to 20 nM for measurements. A significant population of particles corresponding to tetrameric β_2_AR is detected in (**F**).

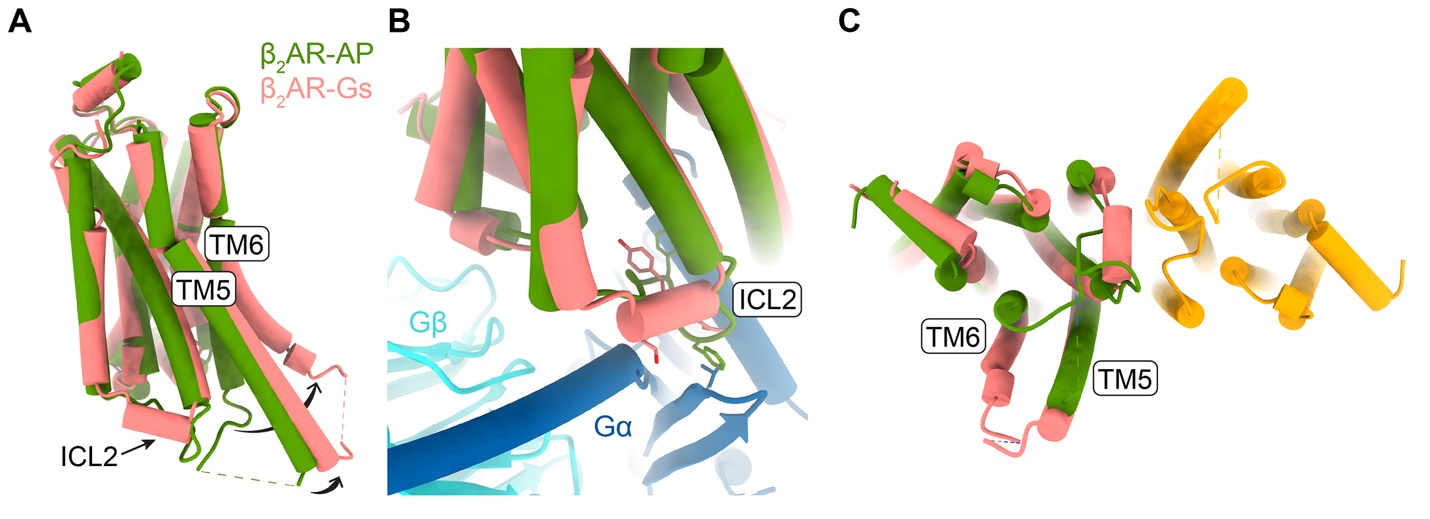

**Figure S9. Conformational changes between AP-bound β₂AR and Gs-bound activated β₂AR.** (**A**) Structural comparison of one protomer from the AP-bound dimer (green) and monomeric Gs-bound β_2_AR (pink, PDB ID: 3SN6). (**B**) Conformational changes in ICL2 in the active β_2_AR. Gs bound to active β_2_AR is shown in dark blue and cyan. Side chains of key contact residues are shown as sticks. (**C**) Intracellular view of active β_2_AR (pink) superimposed onto one protomer (green) of the AP-bound dimer (green and orange).

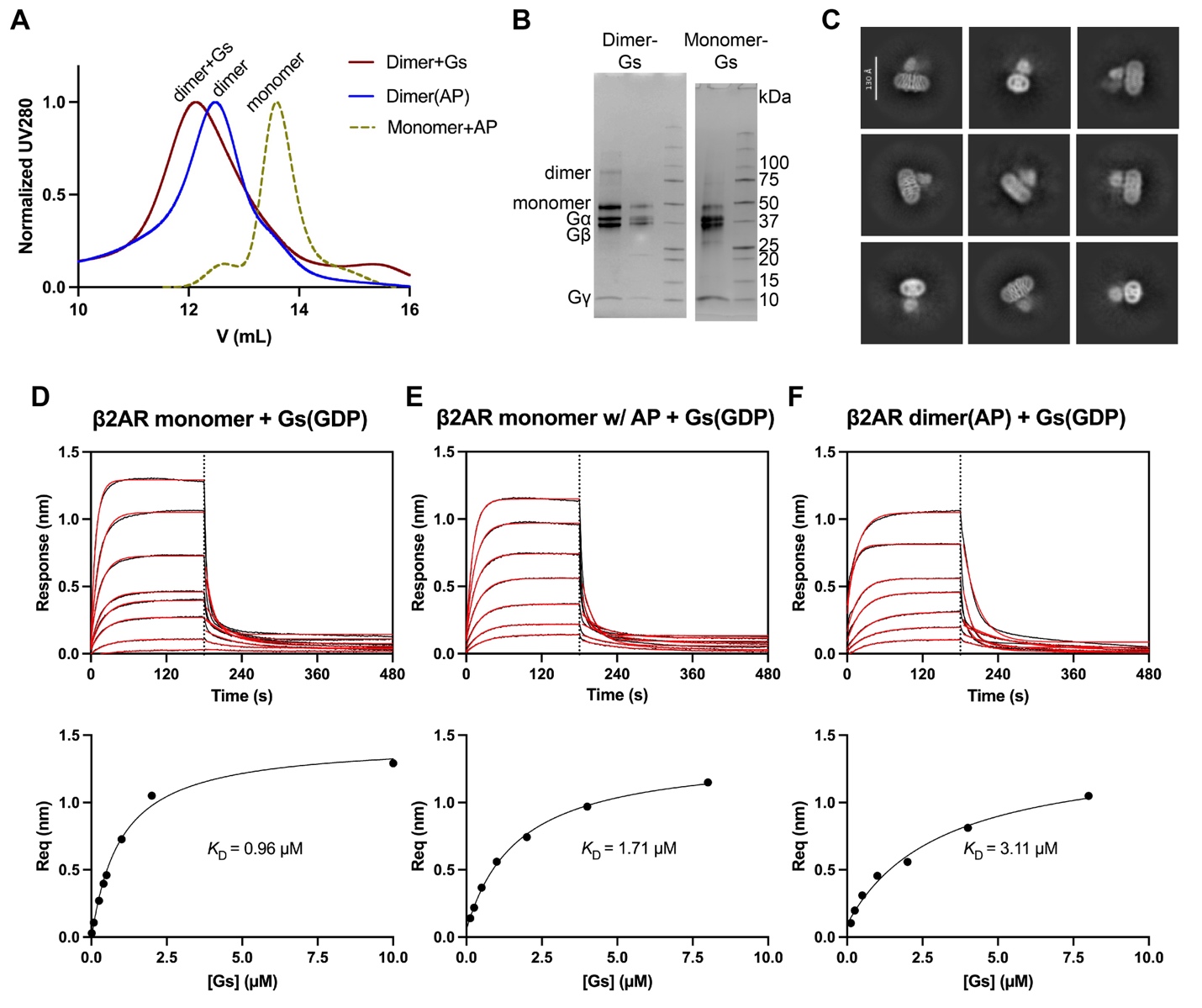

**Figure S10.** **AP-bound β₂AR dimer can effectively form a complex with Gs.** (**A**) SEC profiles showing the formation of a stable dimer–Gs complex after GDP removal by apyrase. (**B**) Coomassie-stained SDS-PAGE gel of peak fractions from (**A**). (**C**) Preliminary cryo-EM 2D class averages of the dimer–Gs complex. Sensorgrams of BLI binding assays of GDP-bound Gs to (**D**) β_2_AR monomer, (**E**) monomer with 10 µM AP, and (**F**) AP-bound β_2_AR dimer. Iso is present at 10 µM in all the conditions. Kinetics and equilibrium binding constants are summarized in **Table S2**.

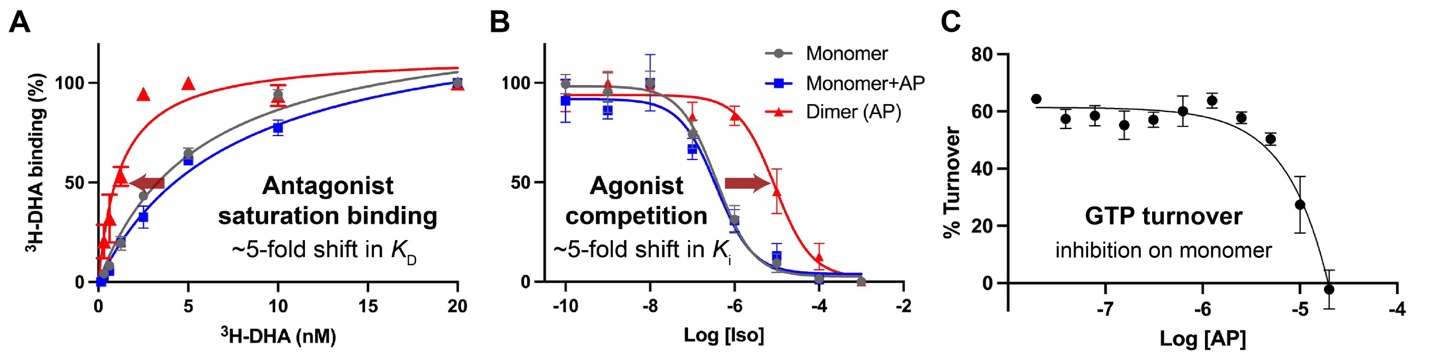

**Figure S11.** **Radioligand binding and GTP turnover assays of β_2_AR in the presence of AP.** (**A**) Saturation binding of β_2_AR to the radiolabeled antagonist [^3^H]-DHA. Shifts in binding affinity (*K*_D_) between monomeric β_2_AR with AP and AP-bound dimer are indicated. (**B**) Competition binding of [^3^H]-DHA with agonist Iso in monomeric β_2_AR (gray), monomer with 10 µM AP (blue), and AP-bound dimer (red). Shifts in the inhibition constant (*K*_i_) of Iso are indicated. (**C**) GTP turnover by Gs, catalyzed by monomeric β_2_AR, is inhibited by AP at high concentrations. All assays were performed with β₂AR reconstituted in nanodiscs. Data are shown as mean with SEM. In (**A**) and (**B**), n = 3; in (**C**), n = 5.

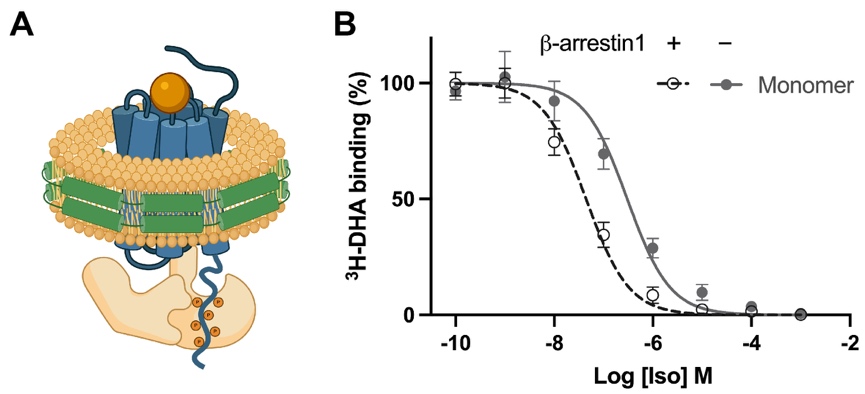

**Figure S12.** **Allosteric effect of β-arrestin1 on radioligand binding of β₂AR.** (**A**) Schematic of β_2_AR ligated to V_2_Rpp in nanodisc illustrating core engagement with β-arrestin. (**B**) Competition binding of [^3^H]-DHA with agonist Iso in monomeric β_2_AR in the presence (black) or absence (gray) of β-arrestin1.

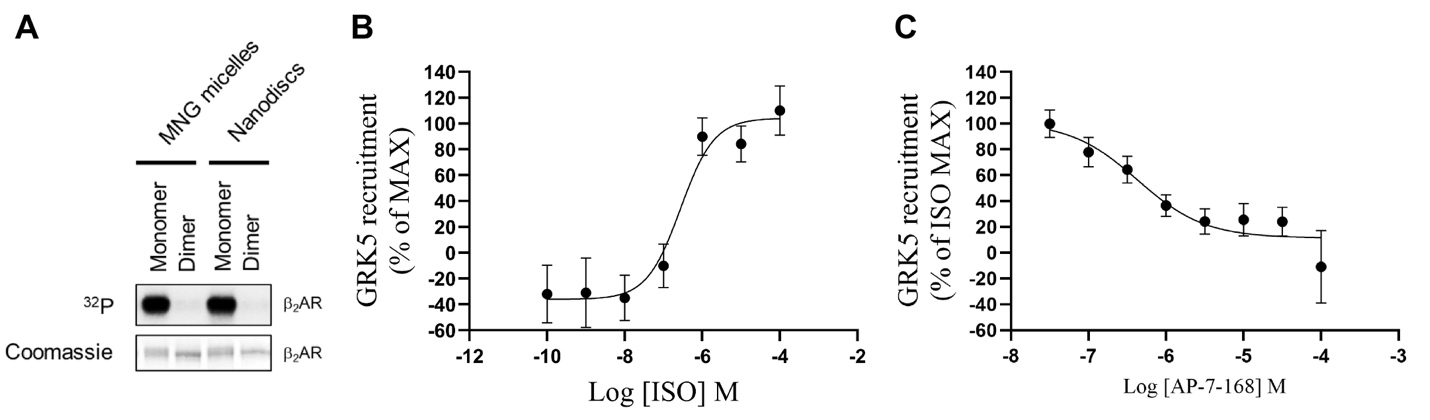

**Figure S13.** **β_2_AR phosphorylation and GRK5 recruitment in cells.** (**A**) Radiometric kinase assay of β_2_AR in detergent or nanodisc. Left: Coomassie staining and ^32^P autoradiography of β_2_AR. Quantification of phosphorylation levels is shown in Fig. 5H. (**B**) Dose-dependent curve of ISO-stimulated GRK5 recruitment in cells. (**C**) Dose-dependent inhibition of GRK5 recruitment by AP. 1 µM ISO was used to stimulate GRK5 recruitment. All data are presented as mean with SEM, n = 3.

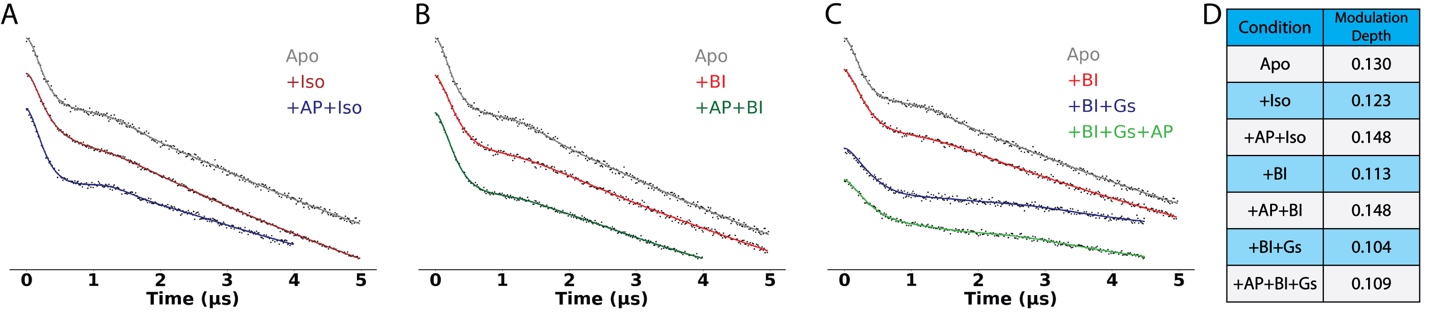

**Figure S14. DEER dipolar evolution traces of spin-labeled β_2_AR in detergent micelles.** DEER measurements were performed on β_2_AR with spin labels attached to TM4 and TM6. **(A–C)** Background-corrected dipolar evolution traces under different ligand conditions. Black dots represent experimental data; colored lines represent fitted curves which are used to derive the distance distributions shown in **Figure 6F–H**. **(D)** The modulation depths for each experimental condition.

**Table S1. Cryo-EM data collection, refinement, and validation statistics**

| **Protein** | **Human β_2_AR_dimer(AP) in MSPE3D1 nanodisc** | **Human β_2_AR_dimer(AP) in detergent** |
| --- | --- | --- |
| **Cryo-EM Data Collection** |  |  |
| Voltage (kV) | 300 | 300 |
| Magnification (x) | 130,000 | 130,000 |
| Pixel Size (Å) | 0.83 | 0.95 |
| Electron exposure (e^-^/Å^2^/frame) | 1.25 | 1.25 |
| Defocus range (µm) | [−2.0, −0.8] | [−2.0, −0.8] |
| Number of image stacks | 17,997 | 7,139 |
| Number of frames per stack | 40 | 40 |
| **Cryo-EM Data Processing** |  |  |
| Initial number of particles | 2,371,430 | 1,961,893 |
| Final number of particles | 237,408 | 48,805 |
| Symmetry imposed | C2 | C2 |
| Map sharpening B factor (Å^2^) | 100 | 125 |
| Map resolution (Å) | 2.5 | 2.9 |
| Map resolution range (Å) | 2.1 – 2.8 | 2.4 – 3.5 |
| FSC threshold | 0.143 | 0.143 |
| **Model Refinement** |  |  |
| Number of amino acids | 287 |  |
| Total non-hydrogen atoms | 2479 |  |
| Model-to-map resolution (Å) | 2.6 |  |
| Average B factor (Å^2^) | 18.10 |  |
| Bond length RMSD (Å) | 0.003 |  |
| Bond angle RMSD (°) | 0.513 |  |
| Ramachandran Plot |  |  |
| Favored (%) | 100 |  |
| Allowed (%) | 0 |  |
| Outliers (%) | 0 |  |
| Rotamer outliers (%) | 0.39 |  |
| MolProbity Score | 0.98 |  |

**Table S2. Kinetic and binding constants for Gs(GDP) interactions with β₂AR measured by BLI**

| Sample | monomer | monomer + AP | dimer (AP) |
| --- | --- | --- | --- |
| *K*_D_ (µM) | 0.96 ± 0.30 | 1.71 ± 0.48 | 3.11 ± 0.89 |
| *k*_on_ (×10^4^ M^-1^ s^-1^) | 7.53 ± 1.53 | 5.37 ± 2.80 | 2.82 ± 2.22 |
| *k*_off_ (×10^-2^ s^-1^) | 2.42 ± 0.22 | 2.30 ± 0.68 | 1.57 ± 0.10 |
